# Substrate specificity in a designed RAS-targeting protease is coupled to active site and distal motions

**DOI:** 10.64898/2026.01.15.699477

**Authors:** Betty Chu, Yanan He, Yihong Chen, Eric A. Toth, John Orban

**Affiliations:** W. M. Keck Laboratory for Structural Biology, University of Maryland Institute for Bioscience and Biotechnology Research, Rockville, Maryland, USA; Department of Chemistry and Biochemistry, University of Maryland, College Park, Maryland, USA

**Author notes:** Corresponding Author(s): John Orban, 9600 Gudelsky Drive, Rockville, MD 20850, Eric A. Toth, 9600 Gudelsky Drive, Rockville, MD 20850.

**Keywords:** RAS, Protease, Dynamic Allostery, Substrate Specificity, Protein Design

## Abstract

Designing proteases with tailored substrate specificity has emerged as a powerful strategy for manipulating protein function in cells. RAS, a key regulator of cell survival and proliferation, is a compelling target for such approaches. Mutations in RAS are involved in about one-third of all human cancers and drive the hyperactive signaling that promotes tumorigenesis, growth, and metastasis in cancers such as pancreatic and lung cancer. This creates a pressing need for strategies capable of modulating mutant RAS with high substrate specificity to avoid unintended cleavage events. As a model for targeted proteolysis, we present the high-resolution crystal structures of RASProtease(II), which provide a detailed view of the enzyme’s active site and substrate-binding architecture. Kinetic experiments showed that cleavage of the cognate QEEYSAM substrate is approximately 30-fold faster than the non-cognate QEEISAM, demonstrating strong proteolytic selectivity. NMR dynamics studies combined with structural mapping revealed that substrate binding modulates not only the active site, but also distal regions of RASProtease(II), uncovering long-range allosteric networks. Contrary to the conventional view that non-cognate substrates are simply poor fits for the active site, we found that binding of the non-cognate peptide induces a greater amount of conformational dynamics in the protease than in the apo form or cognate complex, resulting in significant destabilization and providing a mechanistic explanation for the reduced catalytic efficiency. These results reveal how distal structural networks help define substrate specificity and provide principles for rationally designing proteases with enhanced specificity for therapeutic applications.

**Significance Statement:** Designer proteases are gaining interest for their potential as therapeutic agents. It is therefore critical to understand the mechanisms that differentiate cognate and non-cognate sequences to minimize deleterious off-target cleavage events. While the conventional view is that non-cognate substrates are simply poor fits for the active site, we demonstrate here that dynamic allostery plays an integral role in distinguishing between cognate and off-target sequences, using a protease designed to target the oncoprotein RAS as our model system. Moreover, because the RASProtease is derived from a prototypical serine protease, subtilisin, it is likely that the overall mechanism employed for selecting cognate over non-cognate substrates presented here will be generalizable, providing principles for designing enhanced specificity in a wide range of proteases.

## Introduction

Engineering proteases to conduct site-specific cleavage of proteins that drive disease states has the potential to markedly expand the space accessible to therapeutic intervention. Leveraging the catalytic turnover of proteases could thus facilitate lower dosing requirements and irreversible inactivation of a given target. There are multiple avenues for altering the specificity of proteases to target a potential cleavage site on the target of interest, from structure-based mutagenesis to directed evolution (1). However, once the desired target cleavage is achieved, ensuring high specificity of cleavage becomes the next goal. The process of altering cleavage site preference can at times impair specificity, as was observed for an engineered variant of TEV protease that cleaves the pro-inflammatory cytokine IL-23 (2). TEV protease typically has high specificity, so the observed broadening of specificity aptly highlights the challenge that emerges during the design process. Ideally, a designer protease would have high specificity to avoid deleterious off-target cleavage effects (3). For subtilisin-type proteases, which can have modest specificity for P2 and P3 sites (4–6), the avenues for tightening specificity might not be obvious. Thus, it is important to develop a more comprehensive understanding of what differentiates cognate and non-cognate sequences. Moreover, allostery modulates the activity of proteases and other enzymes, [reviewed in (7–10)] yet the role of allostery in substrate specificity is not well understood. This is especially pertinent to dynamic allostery, which entails ligand-induced alterations of both local and global dynamic motion (8, 11–14) without large-scale conformational shifts, and can regulate enzyme activity via long-range communication, even if the protein is monomeric (15). The effect of cognate versus non-cognate substrates on dynamic allostery communication pathways is relatively unexplored and could open up new reservoirs of residues to be exploited by design efforts.

RAS is a central signaling regulator that controls cell survival and proliferation. As such, mutations in one of the three *Ras* genes (*KRas*, *HRas*, or *NRas*) are involved in roughly a third of all human cancers (16). RAS exists in two states, GTP-bound (active) and GDP-bound (inactive), which are structurally and dynamically distinct. Our previous work (4) showed that, in the active form, the switch I and switch II regions of RAS are more dynamic on the µs–ms timescale than in the inactive form, highlighting a potential vulnerability to protease cleavage. The active state favors cell proliferation, causing oncogenic RAS mutants to be invariably trapped in the activated GTP-bound state, resulting in hyperactivity of signaling pathways that drive tumorigenesis, growth, and metastasis of several cancers. As such, mutant RAS has been a long-standing target for downregulation via small-molecule inhibitors, with recent successes starting with the covalent modification of KRAS-G12C (17–23), followed by the development of KRAS-G12D-specific inhibitors (24, 25) and pan-RAS inhibitors (26–30) that directly target RAS activity. Another strategy for regulating RAS activity is via targeted degradation, which has been explored in multiple ways. First, a proof-in-principle demonstration of targeted RAS degradation using a designed ubiquitin ligase (31) showed decreased pERK levels and cell proliferation. Second, the development of <u>PRO</u>teolysis <u>TA</u>rgeting <u>C</u>himeras (PROTACs) to facilitate proteasome-directed removal of RAS (32) elicited similar effects. Finally, the discovery of the **R**AS/**R**AP1-**s**pecific endo**p**eptidase (RRSP), which cleaves both RAS and RAP1 in the switch I region (33–35), depleted RAS in cells and downregulated RAS signaling. Additionally, RRSP has been used to deplete RAS in cancer cell models, including triple-negative breast cancer and patient-derived xenografts. In both cases, there is a robust decrease in cell viability upon RAS elimination and subsequent inhibition of ERK signaling (36–39).

We recently developed proteases that site-specifically cleave RAS at switch II (4). These engineered RASProteases, which require imidazole or nitrite for catalytic activity, leverage the inherent dynamics of switch II to preferentially cleave active RAS (irrespective of the mutation status), which subsequently destabilizes RAS (4). RASProtease possesses a strong preference for tyrosine at P4 as part of the **<u>Y</u>**SAM sequence in switch II, with a specificity constant that is two orders of magnitude greater than that for the same substrate with isoleucine at P4 (**I**SAM). In the present study, we have conducted an assessment of kinetics, structure, and dynamics of a second-generation RASProtease, hereafter referred to as RASProtease(II), in complex with a cognate QEE**<u>Y</u>**SAM or non-cognate QEE**I**SAM peptide. Remarkably, we observed a pronounced change in dynamic motion for the cognate versus non-cognate complexes, both for the bound peptide and for the protease itself. In particular, our results highlight the importance of dynamic allostery in modulating RASProtease(II) reactivity, and provide insights into the basis for distinguishing between cognate and non-cognate substrates. Moreover, because RASProtease(II) is derived from the prototypical serine protease, subtilisin, the overall mechanism for selection between cognate and non-cognate sequences is likely to be generalizable to many other proteases.

## Results

### Complex Formation of RASProtease(II) with Cognate and Non-cognate Peptides

To understand the structural basis for specificity, we determined the crystal structures of the RASProtease(II) alone and in complex with the QEEYSAM and QEEISAM peptides. As expected, the global structure of apo RASProtease(II) is very similar to the progenitor RASProtease (PDB: 6U9L) (4) with an RMSD of 0.42 Å between Cα atoms.

Differing from the previous generation RASProtease, however, RASProtease(II) was crystallized in the monoclinic P2_1_ space group with unit cell dimensions that accommodate two protein chains in the asymmetric unit. The cognate QEEYSAM-bound and non-cognate QEEISAM-bound RASProtease(II) complexes were crystallized in the same space group as that of the apo form, with two chains in the asymmetric unit and one peptide bound to each protein chain. Complex formation with the peptide substrate did not alter the overall structure of the protease, as demonstrated by the RMSD values between the Cα atoms of the apo protease and those of the cognate and non-cognate complexes, which were 0.31 Å and 0.30 Å, respectively. Contacts between the enzyme and substrate occur throughout all 7 residues of the QEEXSAM peptide, denoted P7 through P1, where X = Y or I (**Figure 1A, B)**. Residue positions are designated according to the Schechter and Berger nomenclature (40), with P7–P1 indicating residues N-terminal to the cleavage site and P1′–P2′ indicating C-terminal positions. Corresponding subsites on RASProtease(II) are denoted S1, S2, etc. No P′ residues are resolved in the present structures.

**Figure 1.**
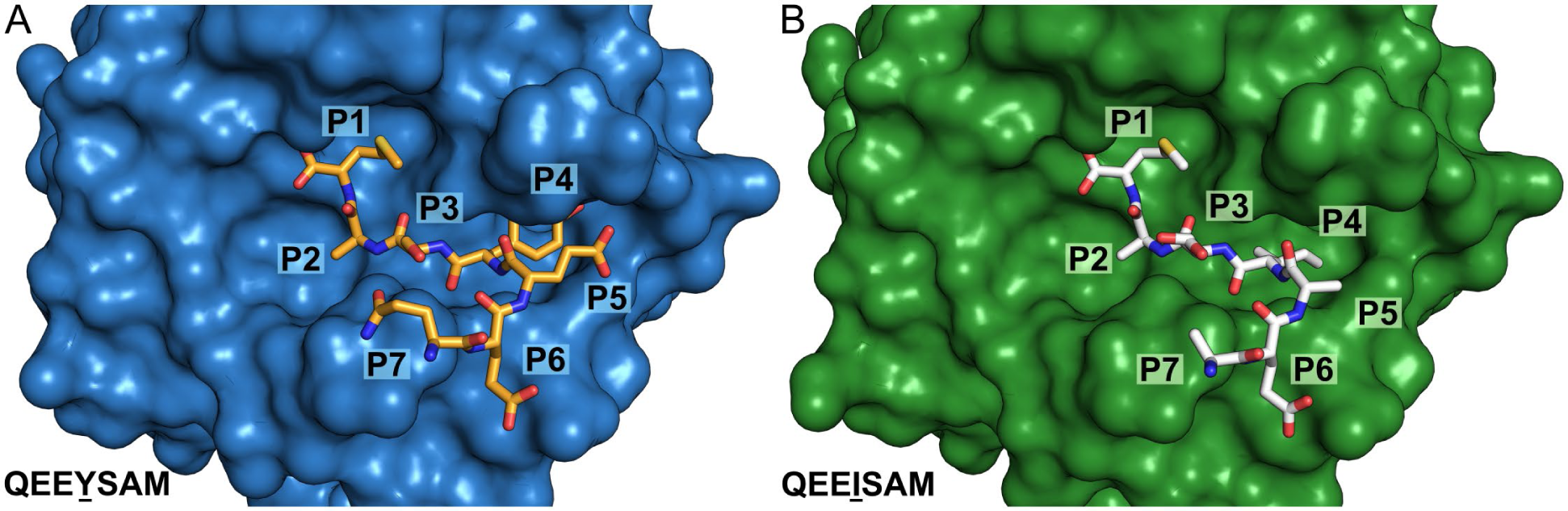
Formation of the protease-peptide complex. Surface representation of RASProtease(II) in complex with the **(A)** cognate QEEYSAM or **(B)** non-cognate QEEISAM peptide. The substrate positions are labeled P1 through P7, where P1 is the cleavage site. Only one protease chain with one bound peptide molecule is shown for clarity.

Electron density for the acyl-enzyme intermediate is visible in the high-resolution crystal structures for both complexes. The S221 Oγ of RASProtease(II) forms an acyl bond with the carbonyl carbon of M7 in the QEEYSAM and QEEISAM substrate **(Figure S1A, B)**. While the high concentrations of crystallization certainly contribute to the visibility of this bond, the presence of the acyl bond is an indication that RASProtease(II) is active and can target both the cognate and non-cognate substrates, as further discussed below.

### Active Site Structural Differences Driven by the P4 Residue

The crystal structure of the cognate complex clearly shows the binding pocket enveloping the tyrosine residue at the P4 position of the QEEYSAM peptide, forming a number of van der Waals interactions to the aromatic ring (**Figure 2A)**. The pocket also contains a water molecule that is tightly bound via hydrogen bonding to the hydroxyl group of the tyrosine (**Figure 2B)**. The tyrosine hydroxyl group also forms interactions with four residues—S128, S130, G131, and V135—at 3.4–3.8 Å distances **(Figure S2)**, further stabilizing the complex. Similarly, the non-cognate complex incorporates the QEEISAM peptide in the binding pocket. With a single amino acid difference at the P4 position, QEEISAM aligns well with its cognate counterpart, showing an RMSD of 0.52 Å for all peptide atoms.

**Figure 2.**
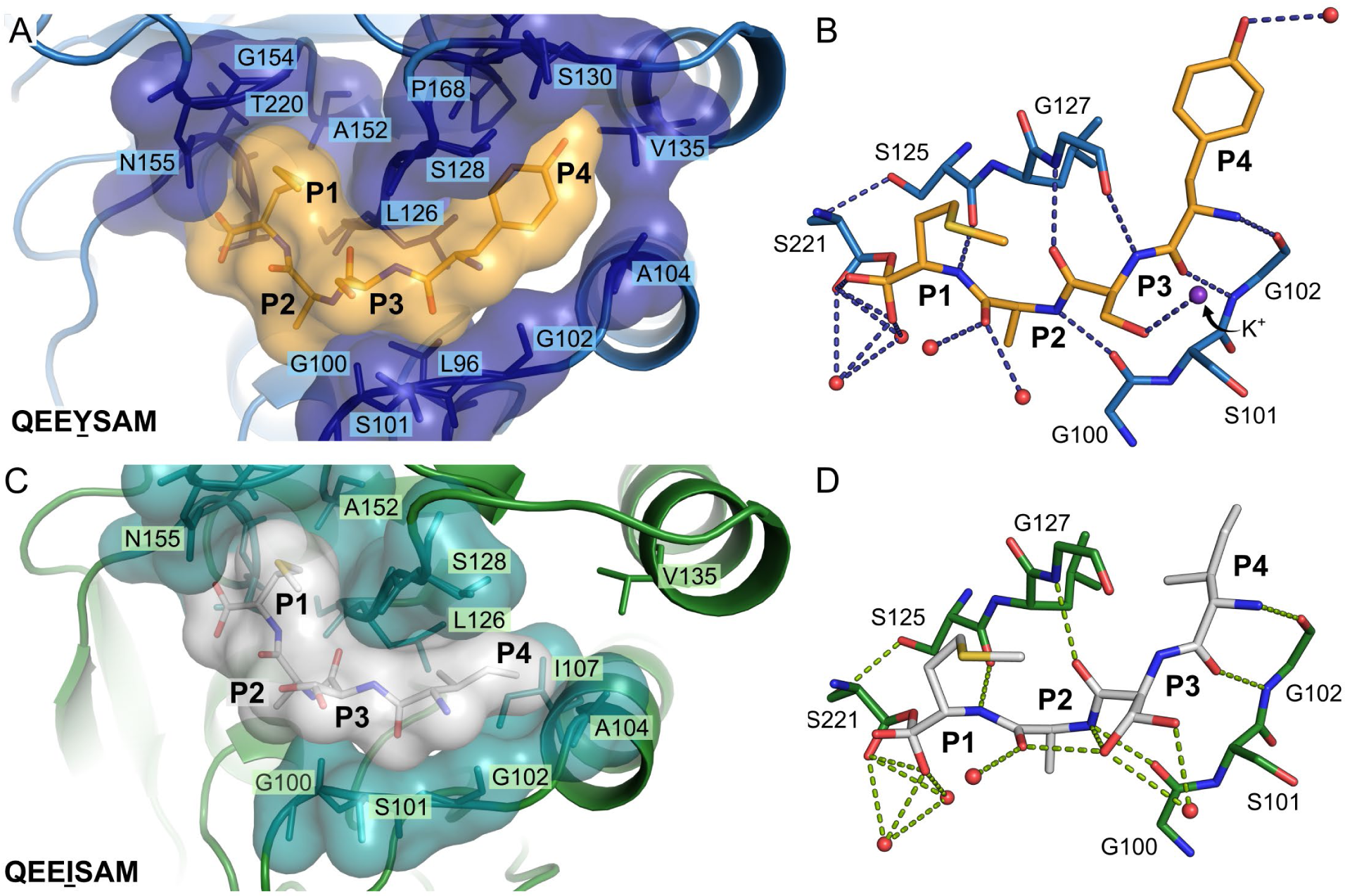
Contacts between RASProtease(II) and the cognate or non-cognate peptides. **(A, B)** Cognate RASProtease(II) is colored dark blue with the peptide atoms colored orange. **(C, D)** Non-cognate RASProtease(II) is colored teal with the peptide atoms colored light gray. **(A, C)** The solvent accessible surface area for RASProtease(II) and its corresponding peptide are displayed to show the van der Waals interaction network that encases the bound peptide. Only P1 through P4 residues of the peptide are shown for clarity. **(B, D)** The hydrogen bonding interactions in the active site are shown as dashes. Waters are represented as red spheres. **(B)** A potassium ion is depicted as a purple sphere in the cognate QEEYSAM complex.

However, the size disparity between the P4 residues, tyrosine and isoleucine, leads to significant structural differences. The large aromatic ring of tyrosine allows it to form stabilizing van der Waals interactions with the alpha helix cap at the C-terminal of residues 133–144. Specifically, the hydroxyl group of tyrosine forms strong interactions with the Cγ_1_ and Cγ_2_ atoms of V135, at distances of 3.7 Å and 3.6 Å, respectively.

Additionally, the tyrosine Cζ atom is positioned within 4.4 Å of the Cγ atom of V135, further contributing to the stability of the local structure. In contrast, the smaller isoleucine side chain is more than 5.0 Å away from the alpha helix cap and is unable to interact with V135, resulting in a loss of van der Waals interactions (**Figure 2C)**. The stability of the non-cognate complex is further reduced by the lack of hydrogen bonding between the nonpolar isoleucine and the water molecules in the binding pocket (**Figure 2D)**. Interestingly, the P4 isoleucine is positioned within 4.0 Å of I107, resulting in a gain of some stabilizing forces, but this does not offset the loss of interactions with the C-terminal alpha helix cap.

### Anisotropic Motion in Binding Pocket and Distal Sites

To gain a deeper understanding of the differences between the cognate and non-cognate complexes differing at P4, we analyzed the anisotropic motion observed in the bound peptides. Because the crystal structures were resolved at high resolution (1.2–1.5 Å), we were able to refine the models using anisotropic displacement parameters, which allowed us to visualize individual atomic movement. The cognate QEEYSAM peptide shows a modest amount of anisotropic motion, with the spheres surrounding the P4 atoms indicating uniform movement (**Figure 3A)**. In contrast, the QEEISAM peptide in the non-cognate complex has significant anisotropic motion. The atoms, particularly those at the P4 position, move in preferred directions, as indicated by the surrounding ellipsoids (**Figure 3B)**.

**Figure 3.**
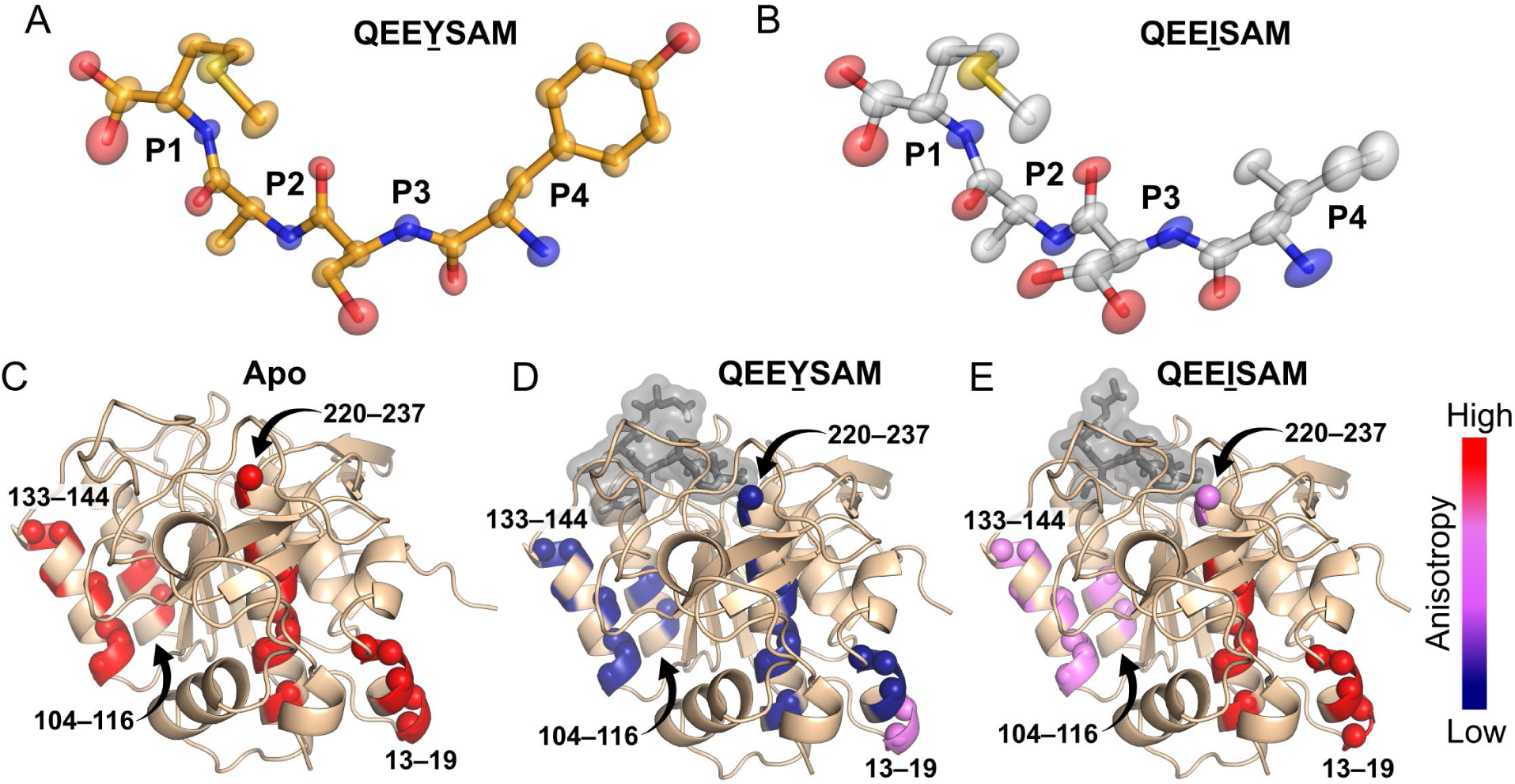
Anisotropic displacement in protease helices and bound peptides in RASProtease(II) structures. The magnitudes and directions of atomic displacement are represented as anisotropic thermal ellipsoids for the bound peptide in **(A)** the cognate QEEYSAM complex and **(B)** the non-cognate QEEISAM complex. Only P1 through P4 residues of the peptide are shown for clarity. **(C)** The most anisotropic residues in α-helices of the apo RASProtease(II) structure were identified, numbered by their residue ranges, and colored according to a continuous scale from blue (low anisotropy) to red (high anisotropy). The same color scale was applied to **(D)** the QEEYSAM-bound and **(E)** the QEEISAM-bound complexes to visualize changes in anisotropy upon peptide binding. In the peptide-bound structures, the peptide is shown as gray sticks with a semi-transparent surface to highlight the active site.

In addition to local anisotropic motion at the active site, peptide binding surprisingly induces significant changes in regions distal to the binding pocket. To examine these effects, we analyzed the anisotropy of each residue of RASProtease(II) in the apo, cognate, and non-cognate complex structures **(Figure S3)**. In the apo RASProtease(II) structure, we identified four helical regions—residues 13–19, 104–116, 133–144, and 220–237—with pronounced anisotropic motion, highlighted in red in **Figure 3C**. Binding of the cognate QEEYSAM peptide dampens anisotropic motion, as indicated by a shift toward blue in the anisotropy color scale in **Figure 3D**. In particular, helices spanning residues 104–116, 133–144, and 220–237 show more isotropic motion upon peptide binding. Interestingly, residues 18 and 19, colored violet in **Figure 3D**, remain anisotropic in the complex, suggesting that the cognate peptide has a relatively small effect on this region. Despite being located distal from the active site, these residues show a change in anisotropy, highlighting distinct effects of cognate peptide binding on the protease structure.

In contrast, binding of the non-cognate QEEISAM peptide results in anisotropy patterns similar to those of the apo form. Helices 104–116 and 133–144 show modest stabilization, as indicated by a shift toward violet in the anisotropy color scale in **Figure 3E**. However, helices 13–19 and 220–237 remain anisotropic with no appreciable change relative to the apo structure, suggesting that the non-cognate complex retains a higher degree of structural flexibility.

### Substrate Specificity of RASProtease(II) for Cognate and Non-cognate Peptides

To assess the activity of RASProtease(II) and to determine the rate of a single catalytic cycle, we collected fluorescence progress curves under single-turnover conditions, in which enzyme concentration is in excess over substrate concentration. The progress curves were fit to Equation (1), allowing for the determination of the observed rate constants (*k*_obs_). We observed a burst phase in the cleavage reaction between RASProtease(II) and the cognate QEEYSAM-AMC substrate (**Figure 4)**. At 4 µM of the enzyme, the *k*_obs_ was determined to be 0.0153770 s^-1^. In contrast, the reaction with the non-cognate QEEISAM-AMC did not exhibit a burst phase, but instead showed a linear increase in fluorescence. A linear fit of the data was used to extract the *k*_obs_, which is directly proportional to the slope. The resulting *k*_obs_ value of 0.0005565 s^-1^ is significantly slower (∼30-fold) than that observed with the cognate substrate. These results underscore the substrate specificity of RASProtease(II) and highlight its strong preference for the cognate peptide.

**Figure 4.**
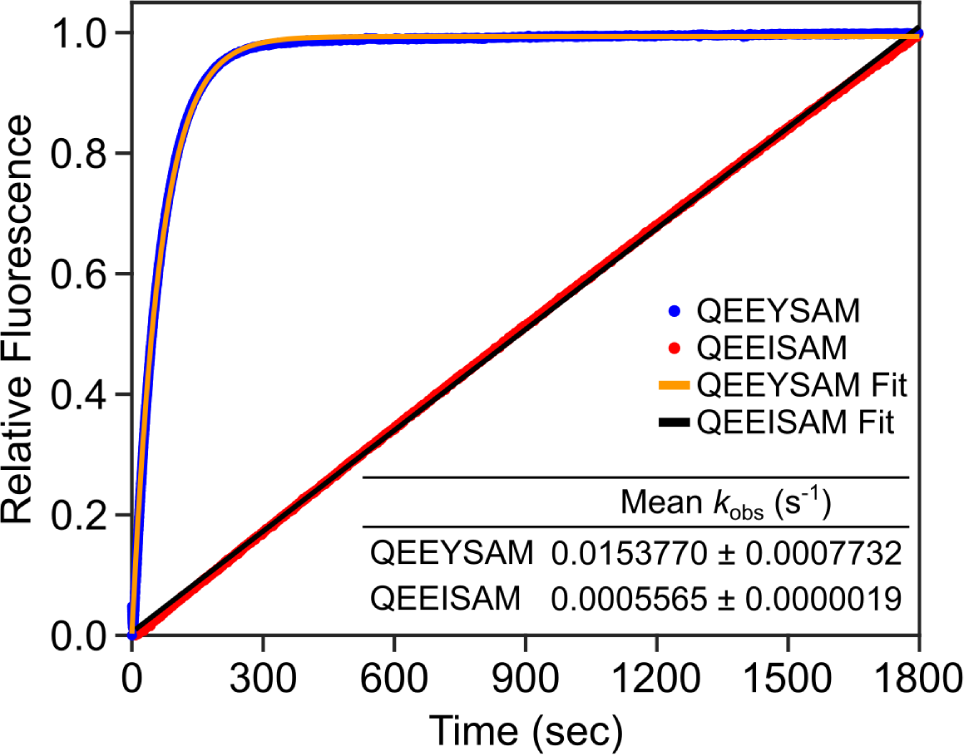
Cleavage of 100 nM cognate QEEYSAM-AMC or non-cognate QEEISAM-AMC peptide by 4 µM RASProtease(II). Representative single-turnover kinetic fluorescent progress curves are shown, highlighting differences in the mean *k*_obs_ values. For the reaction with the non-cognate peptide, the initial 8 seconds of signal contained electronic noise and was omitted for clarity. The cognate and non-cognate progress curves are shown in blue and red, respectively, with the corresponding curve fits overlaid in light orange and black, respectively.

### NMR Analysis of RASProtease(II) Complexes

We undertook a detailed NMR analysis of RASProtease(II) in the apo and peptide-bound states to evaluate the effects of binding in solution and to enable a comparison with the X-ray crystallographic data. First, backbone resonances were assigned for the apo (77%) **(Figure S4)**, QEEYSAM-bound (84%) **(Figure S5)**, and QEEISAM-bound (81%) **(Figure S6)** forms of RASProtease(II). We then investigated the chemical shift perturbations (CSPs) and peak intensity changes in RASProtease(II) upon addition of the cognate QEEYSAM or non-cognate QEEISAM peptide and subsequently mapped the observed effects onto the corresponding crystal structures for visualization.

The addition of QEEYSAM led to significant CSPs in the 2D ^1^H-^15^N TROSY-HSQC spectrum of the protease, with many of the affected residues proximal to the active site (<10 Å from a peptide Cα) as would be expected (**Figure 5A, B)**. Distal CSPs (>10 Å from a peptide Cα) were also observed, particularly in the interior helix (66–84) and adjacent β-hairpin (205–217). Moreover, 22 new backbone amide peaks were detected in the presence of QEEYSAM, presumably due to stabilization of the protease structure. Of these, 8 are proximal and 14 are distal to the bound peptide. In contrast, addition of QEEISAM yielded considerably fewer CSPs in the protease and only 8 (1 proximal, 7 distal) new backbone amide peaks (**Figure 5C, D)**.

**Figure 5.**
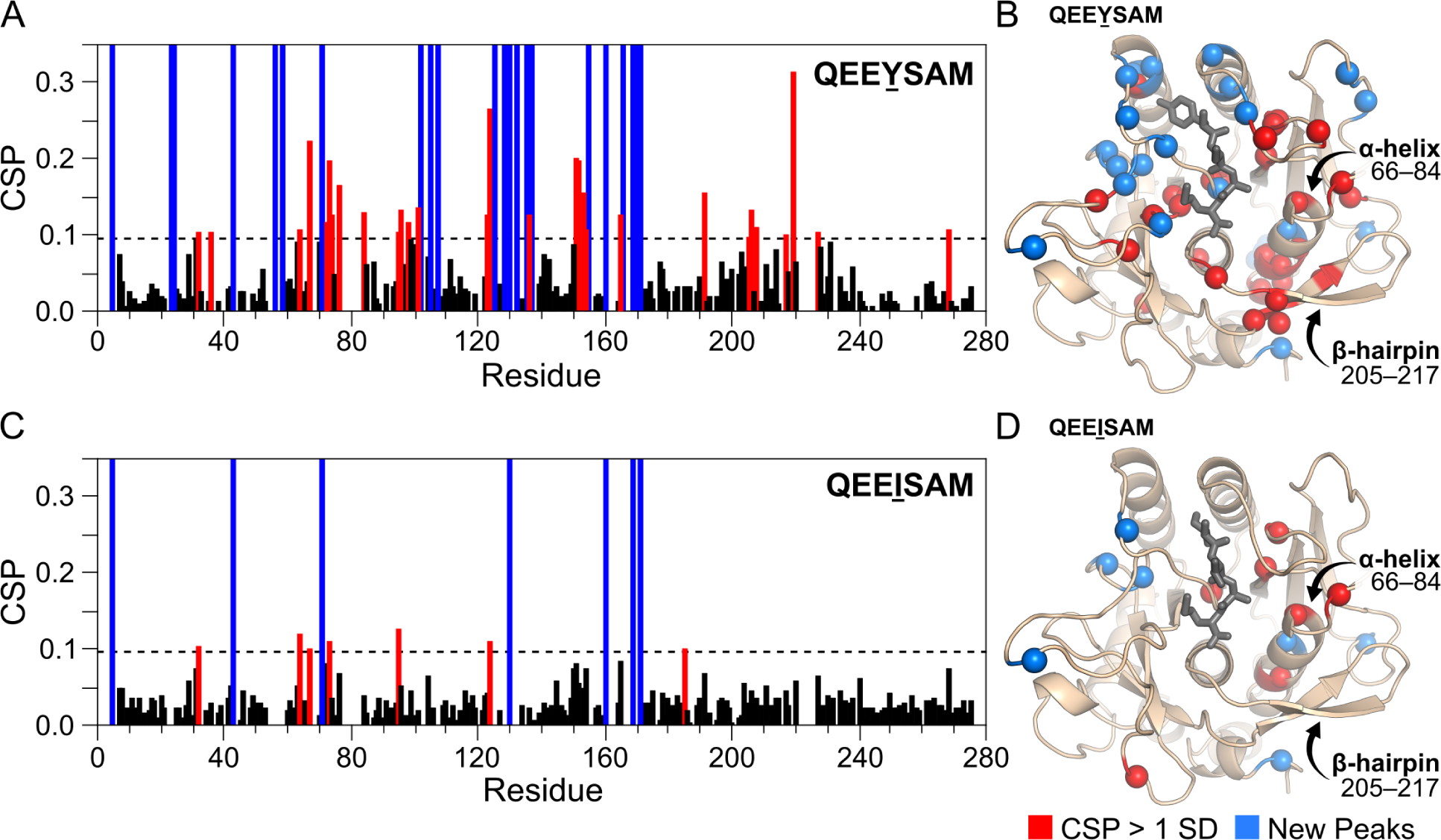
Chemical shift perturbation (CSP) and peak intensity changes in RASProtease(II) upon addition of cognate or non-cognate peptides. **(A, C)** CSPs and new peak appearances for RASProtease(II) upon addition of **(A)** the cognate QEEYSAM peptide or **(C)** the non-cognate QEEISAM peptide are plotted. CSPs are shown as black bars, with the mean ± 1 SD (calculated from the QEEYSAM dataset) indicated by a dashed line. This same cutoff was applied to the QEEISAM data for comparison. Residues with CSPs exceeding this threshold are shown as red bars. New peaks observed upon peptide addition are shown as blue bars. **(B, D)** Residues with significant CSPs (red spheres; CSPs > mean ± 1 SD) and new peaks (blue spheres) are mapped onto the crystal structures of RASProtease(II) in complex with the **(B)** QEEYSAM or **(D)** QEEISAM peptide. Only P1 through P4 residues of the peptide are shown for clarity.

### Fast Timescale Backbone Dynamics

To investigate how cognate and non-cognate substrate binding affects fast (ps–ns) backbone dynamics of RASProtease(II), we measured ^15^N-relaxation parameters (*R*_1_, *R*_2_) and {^1^H}-^15^N steady-state heteronuclear NOEs (hetNOEs) for the apo and peptide-bound states. In all three states, backbone regions exhibited generally high hetNOE values that are consistent with ordered structure **(Figure S7)**. Analysis of hetNOE differences (ΔhetNOE) between the apo and each peptide state, however, showed that overall changes were modest (0.1–0.2), while residue-specific analysis revealed distinct, peptide-dependent changes in backbone flexibility **(Figure S8A, B)**. Addition of the cognate QEEYSAM peptide reduced mobility in residues D99, G100, and A104, adjacent to the S2 and S4 pockets of the active site **(Figure S8C)**, whereas binding of the non-cognate QEEISAM peptide increased flexibility in residues M124 and A151, near the S4 pocket **(Figure S8D)**. These observations indicate that substrate identity differentially modulates protease backbone dynamics, which are further supported by subsequent relaxation measurements.

More significant differences were observed in the individual *R*_1_ and *R*_2_ values between the apo and peptide-bound states (**Figure S9A, B**). Analysis of *R*_2_/*R*_1_ ratios provided insight into the extent of conformational exchange in each state **(Figure S10A)**. Residues with *R*_2_/*R*_1_ values beyond ±1 standard deviation from the mean were identified as positions with substantial backbone mobility. The apo and QEEISAM-bound states exhibited broader *R*_2_/*R*_1_ distributions and a similar number of outlier residues **(Figure S10B, C)**, whereas the QEEYSAM-bound state displayed tightly clustered values and markedly fewer outliers **(Figure S10D)**. This pattern is consistent with reduced conformational exchange and stabilization of backbone dynamics upon cognate peptide binding.

### Exchange-Broadened Signals in RASProtease(II)

Consistent with the above observations suggesting conformational exchange, the apo state has the largest number of undetected backbone amide signals (52 residues), followed by the QEEISAM-bound state (45 residues) and QEEYSAM-bound state (30 residues). The NMR signals corresponding to these residues are broadened beyond detection, and are referred to here as putatively exchange-broadened. Many of these residues occur in solvent accessible loop regions, as is commonly observed in other proteins.

However, a distinct subset of putatively exchange-broadened residues was also observed near the active site and S1–S4 regions of RASProtease(II). Of this subset, 13 residues are common to the apo, QEEISAM-bound, and QEEYSAM-bound states **(Figure S11)**. Several of these residues are located in loop regions adjacent to the docking peptide, including Q103, G131, and Y167 near the S4 pocket, L126 near the S3 site, and S156 near S1, suggesting local flexibility at key peptide-binding interfaces. The exterior helix containing A133 also lies near the S4 pocket, highlighting additional dynamic elements on the protein surface.

Extending from the surface into the protein interior, we observed a group of exchange-broadened residues located in buried helices at the catalytic core. Residues T68 and H69 are part of an interior helix (66–84) and are near the active site residue G67 (mutated from H67 but rescued with the cofactor imidazole (4)). Most notably, the catalytic S221 and adjacent residues M222, A223, S224, and H226 in the central helix (220–237) are also exchange-broadened, reflecting dynamic fluctuations within the protein core. The broadening of these signals indicates that these buried regions participate in conformational exchange processes that are likely to play a critical role in RASProtease(II) catalysis.

### Allosteric Networks Revealed by Relaxation Dispersion

To further quantify the contributions of conformational exchange at the residue level, we employed CPMG relaxation dispersion (RD) experiments, which probe intermediate timescale (µs–ms) dynamic events. Backbone amide conformational exchange rates (*R*_ex_) were extracted from these data for the apo, cognate-bound, and non-cognate-bound forms of RASProtease(II) to assess whether these dynamics correlate with the observed differences in catalytic efficiency between the cognate and non-cognate complexes. For the apo state, in addition to the amino acid positions with completely exchange-broadened signals, there were also a number of residues near the active site that have elevated RD profiles with *R*_ex_ values greater than 10 s^-1^ **(Figure S12, black; S13A, B).** These include active site residues D32 (S2) and G67 (S2), as well as residues L96, D99, M124, A152, G219, and T220, all of which line the S1‒S4 pockets that engage the docking peptide. However, conformational exchange was also observed for residues located more than 10 Å from the active site, such as D41, V46, and A151 **(Figure S14)**. Together, these results indicate that the apo form of RASProtease(II) undergoes widespread conformational exchange spanning the active site, peptide-docking regions, and distal parts of the protein.

Upon addition of the cognate QEEYSAM peptide, significant quenching of exchange is observed in the aforementioned residues at the peptide-docking regions **(Figure S12, blue, Figure S13C)**. This reduction is reflected in *R*_ex_ differences (Δ*R*_ex_) between the apo and QEEYSAM-bound states, where negative Δ*R*_ex_ values indicate decreased conformational exchange (**Figure 6A, C)**. Moreover, of the 22 new amide resonances that appear upon cognate peptide binding, 8 are due to proximal residues (G102, S105, I107, S125, S128, S130, S132, and N155), further indicating a decrease in conformational exchange in the active site. Other amino acid positions, however, exhibit positive Δ*R*_ex_ values (Δ*R*_ex_ 5–10 s^-1^), which correspond to increased conformational exchange. Notably, several of these residues form three distinct clusters with inter-residue heavy atom distances within 5 Å, denoted as (i) N64-D62-I35-L90-A45-S89-C87, (ii) K136-V139-S173-V174-I175-V177-A153-T164-S159, and (iii) T73-V84-H17-Q19-K237-A272-A274-W241-V246 (**Figure 7A)**. Most of these residues are distal to the active site with the exception of N64 in (i) and A153 in (ii), which are located near the S2 and S1 pockets, respectively.

**Figure 6.**
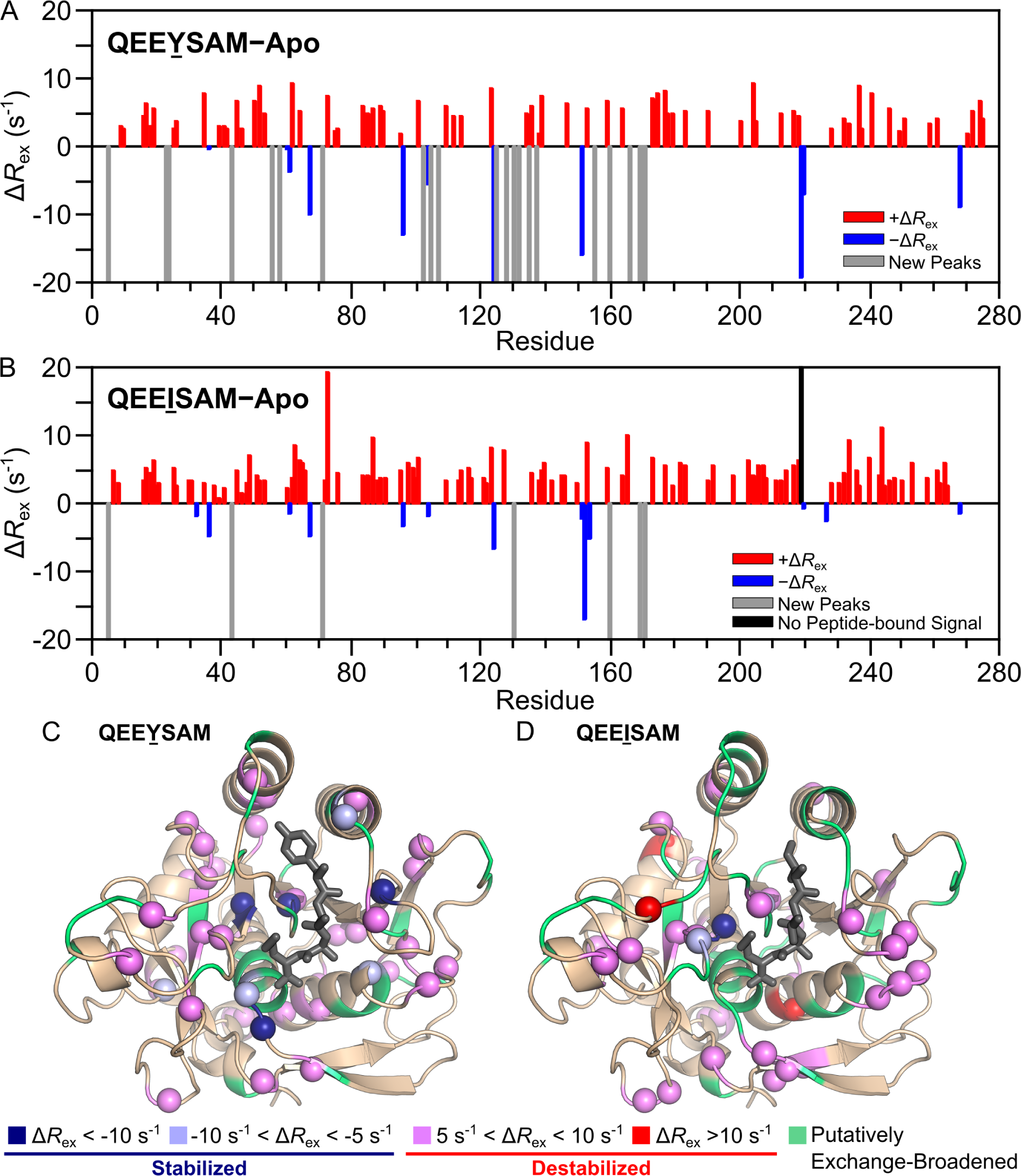
Differences in conformational exchange dynamics between the RASProtease(II) cognate and non-cognate complexes. Residue-specific differences in conformational exchange contributions (Δ*R*_ex_) between the apo and peptide-bound forms were calculated separately for the QEEYSAM and QEEISAM complexes. The Δ*R*_ex_ plots show the difference between **(A)** apo and QEEYSAM and between **(B)** apo and QEEISAM. Positive Δ*R*_ex_ values (red bars) indicate increased conformational exchange upon peptide binding, whereas negative Δ*R*_ex_ values (blue bars) indicate decreased exchange. Gray bars indicate residues where signal appeared upon peptide binding. The black bar represents residue G219, where signal was present in the apo form but missing in the QEEISAM-bound form. **(C, D)** Residues with significant conformational exchange differences are mapped onto the corresponding crystal structures of the **(C)** QEEYSAM-bound and **(D)** QEEISAM-bound complexes. These residues are displayed as spheres and colored according to Δ*R*_ex_ magnitude: dark blue (Δ*R*_ex_ < −10 s^-1^), light purple (−10 s^-1^ < Δ*R*_ex_ < −5 s^-1^), violet (5 s^-1^ < Δ*R*_ex_ < 10 s^-1^), and red (Δ*R*_ex_ > 10 s^-1^). Residues lacking NMR assignments are presumed to be exchange-broadened and are collectively labeled “Putatively Exchange-Broadened” (light green). Only P1 through P4 residues of the peptide are shown for clarity.

**Figure 7.**
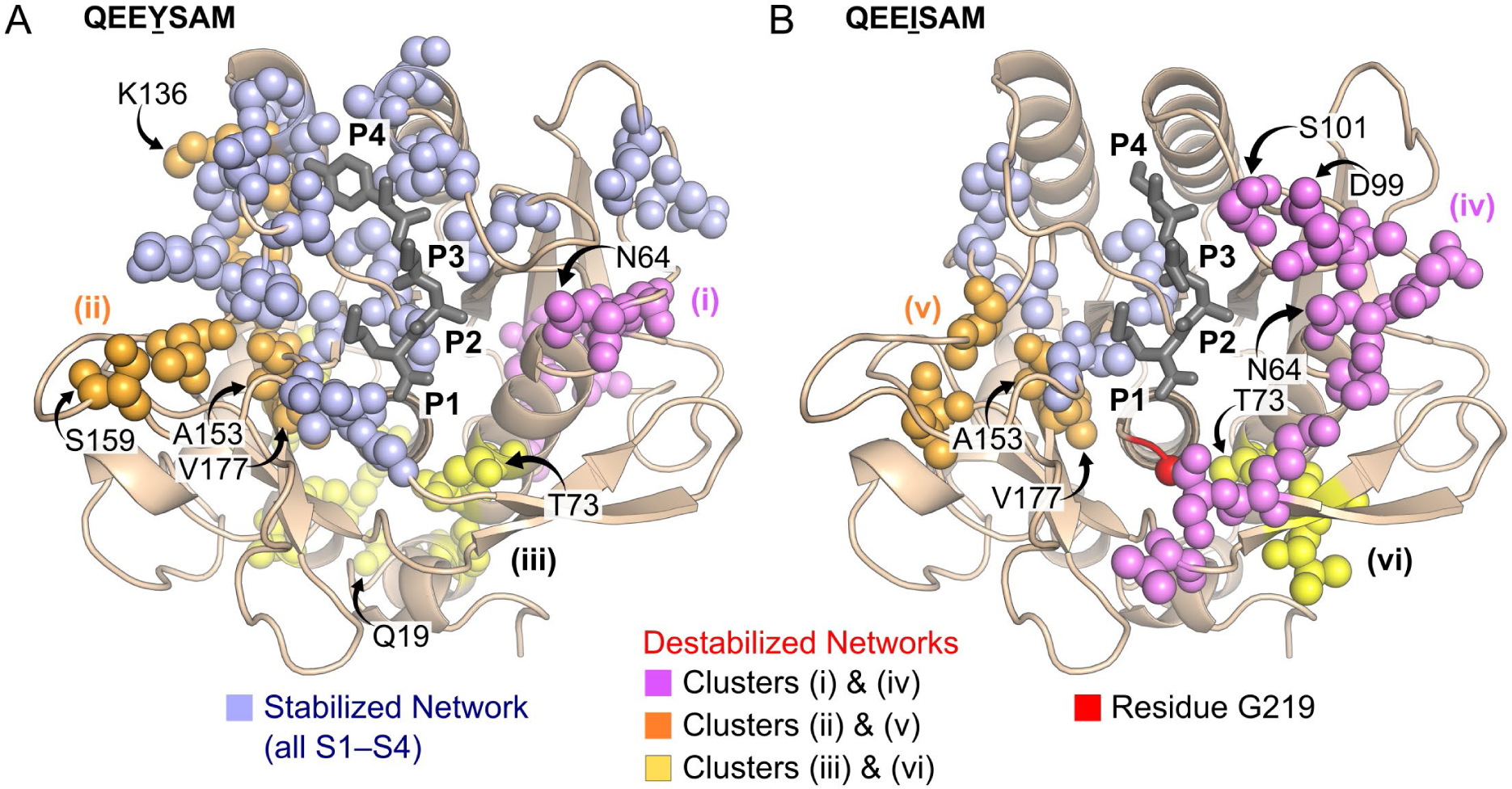
Comparison of allosteric networks identified in the cognate QEEYSAM and non-cognate RASProtease(II) complexes. Analysis of conformational exchange differences (Δ*R*_ex_) between the apo and peptide-bound states revealed residues forming networks with inter-residue heavy atom distances < 5 Å that influence complex stability. These residues are shown as spheres mapped onto the crystal structures of the **(A)** QEEYSAM and **(B)** QEEISAM complexes. Only P1 through P4 residues of the peptide are shown for clarity. Residues with negative Δ*R*_ex_ values (< −5 s^-1^) and within 5 Å of each other form stabilizing networks (light purple), particularly near the S1–S4 substrate-binding pockets. This includes residues for which near peaks appear upon addition of the peptide. Residues with positive Δ*R*_ex_ values (> 5 s^-1^) and within 5 Å of each other contribute to three destabilizing networks: clusters (i) and (iv) (violet), clusters (ii) and (v) (orange), and clusters (iii) and (vi) (yellow). Residue G219, which exchange-broadens beyond detection in the non-cognate complex, is shown as a red sphere to highlight its position in the S1 site.

In contrast, the pattern of conformational exchange in the RASProtease(II) complex with the non-cognate QEEISAM peptide differs significantly. While there is some quenching of exchange in the active site region, it is a smaller effect involving substantially fewer residues than with the cognate peptide (**Figure 6B, D; Figure S13D)**. Only 1 new amide peak (S130) is detected in the proximal region of the non-cognate complex, while the other 7 active site resonances that appear upon cognate complex formation remain exchange-broadened with the addition of QEEISAM. Further, a significant number of residues around the active site display *increased* conformational exchange relative to the apo state. A contiguous network of residues with positive Δ*R*_ex_ values (Δ*R*_ex_ 5–10 s^-1^) extends from the S3 pocket to beyond the peptide cleavage site, forming cluster (iv) S101-D99-A98-N64-N63-S65-K217-S218-V203 (**Figure 7B)**.

Residue G219, which is at the N-terminus of the central helix (220–237) and near the cleavage site, further contributes to this network by becoming completely exchange-broadened, with its signal undetectable upon addition of QEEISAM, whereas its conformational exchange is substantially quenched in the presence of QEEYSAM **(Figure S12)**. Additionally, the non-cognate complex contains two smaller clusters with increased conformational exchange: cluster (v) V177-A153-V165-V192 (Δ*R*_ex_ 5–10 s^-1^), which contains one proximal residue (A153, near S1), and cluster (vi) T73-S207-V206 (Δ*R*_ex_ 5–19 s^-1^), which is entirely distal (**Figure 7B)**. Collectively, the CPMG relaxation analysis reveals clear differences between the conformational exchange profiles of the cognate and non-cognate forms of the RASProtease(II).

### Peptide-Dependent Conformational Exchange Characterized by CEST

We also employed chemical exchange saturation transfer (CEST) experiments to probe exchange processes on an overlapping but slower timescale (∼100 µs to ∼10 ms). CEST analysis showed profiles that varied with the binding state of RASProtease(II), mirroring the conformational exchange patterns observed in the RD study. In the apo state, residues proximal to the S1–S4 pockets exhibited broadened CEST profiles, indicative of exchange between multiple conformational states (**Figure 8A; Figure S15, black)**. Addition of the cognate QEEYSAM peptide significantly narrowed all 13 of these profiles, suggesting structural stabilization and a shift toward a population of more homogeneous conformational ensembles (**Figure 8A; Figure S15, blue)**. In contrast, addition of the non-cognate QEEISAM peptide to RASProtease(II) had little effect on the CEST profiles of the S1–S4 residues. Only 7 of the 13 proximal residues displayed narrowed profiles, while the rest were similar to or even broader than those observed in the apo form, indicating substantial conformational exchange (**Figure 8A; Figure S15, red)**. In particular, residue G219 showed a broad CEST profile in the apo state that narrowed considerably in the presence of the cognate QEEYSAM peptide, but was completely unmeasurable in the non-cognate QEEISAM complex, suggesting significant exchange-broadening **(Figure S15)**.

**Figure 8.**
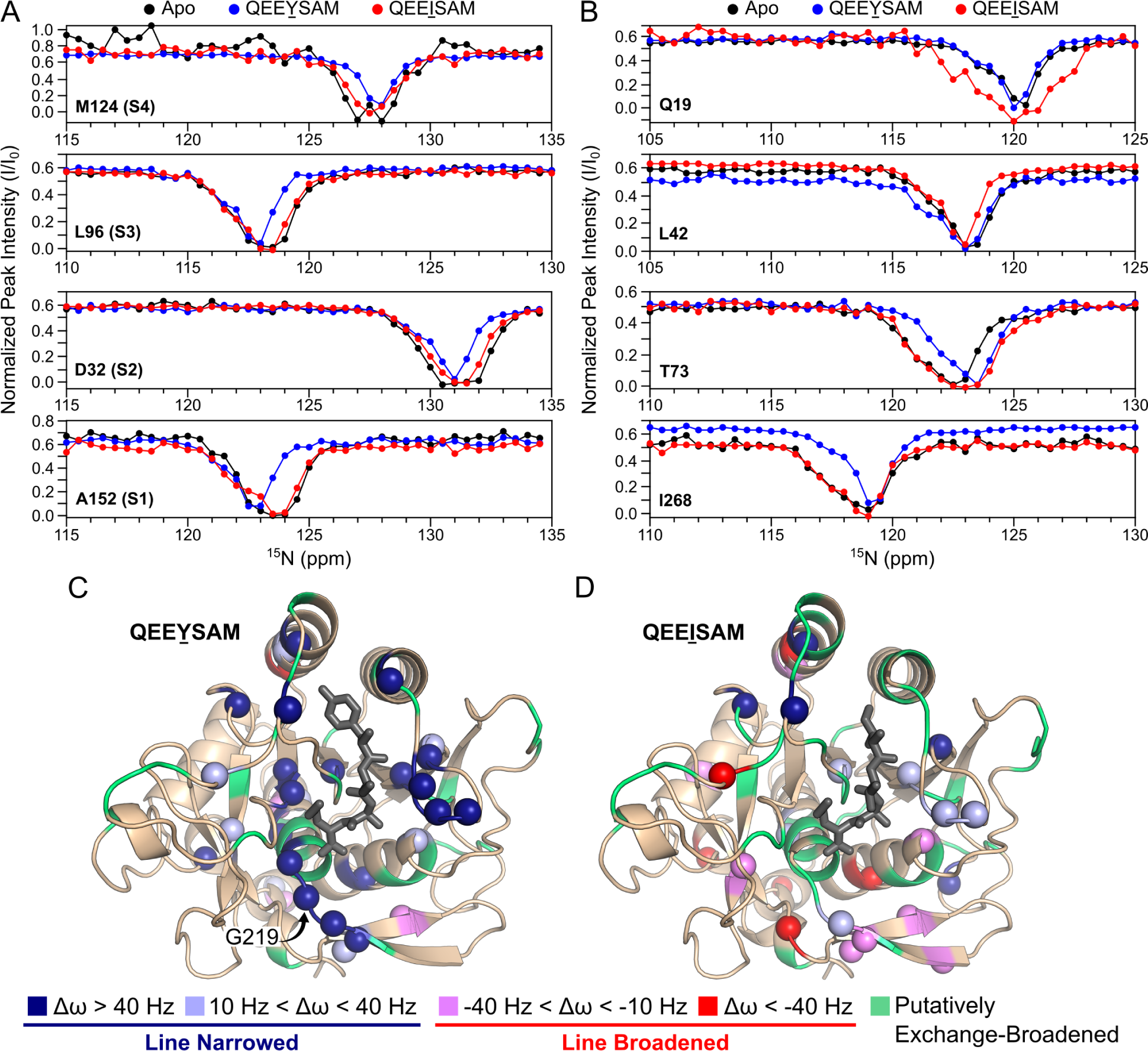
Differences in residue dynamics among the apo, QEEYSAM-bound, and QEEISAM-bound forms of RASProtease(II) revealed by chemical exchange saturation transfer (CEST) experiments. CEST profiles are shown for **(A)** four proximal (M124, L96, D32, A152) and **(B)** four distal (Q19, L42, T73, I268) residues. For proximal residues, the nearest active site pocket (S1–S4) is noted in parentheses. CEST data are presented for the apo form (black), cognate QEEYSAM complex (blue), and non-cognate QEEISAM complex (red), revealing changes in exchange behavior upon peptide binding. **(C, D)** Residues with significant CEST profile differences (Δω, Hz) are mapped onto the crystal structures of the **(C)** QEEYSAM-bound and **(D)** QEEISAM-bound complexes. These residues are shown as spheres and colored by Δω magnitudes as follows: dark blue (Δω > 40 Hz), light purple (10 Hz < Δω < 40 Hz), violet (−40 Hz < Δω < −10 Hz), and red (Δω < −40 Hz). Residues with positive Δω values are line narrowed, whereas those with negative Δω values are line broadened. Residues lacking NMR assignments are presumed to be exchange-broadened and are collectively labeled “Putatively Exchange-Broadened” (light green). Only P1 through P4 residues of the peptide are shown for clarity.

Notably, this effect of increased conformational exchange in the non-cognate complex, but significant quenching in the cognate complex not only occurs across the four substrate-binding pockets (S1–S4) of RASProtease(II), but also in distal regions of the protease. Analysis of residue-specific CEST profile differences (Δω) between the apo and peptide-bound states quantified the extent of conformational exchange associated with peptide binding. In the cognate complex, 31 residues exhibited positive Δω values, reflecting reduced conformational exchange in both the active site and overall structure (**Figure 8C; Figure S16A)**. Interestingly, we noted 6 distal residues (S18, L42, V143, T208, A231, A272) with negative Δω values, which correspond to increased conformational exchange and corroborate the broadened CEST profiles (**Figure 8B; Figure S17)**. In the non-cognate complex, exchange-broadening effects were more widespread, affecting both proximal and distal regions of RASProtease(II). Only 14 residues showed positive Δω values, indicating a smaller number of stabilized residues compared to the cognate complex. Conversely, 18 residues exhibited negative Δω values, all of which had broadened CEST profiles (**Figure 8D; Figure S16B)**. Of these residues, 16 were distal, including a subset of 5 residues (T73, V165, V203, V206, K217) that appear to participate in destabilizing networks that propagate dynamic effects to more distant parts of the protease **(Figure S17)**. Together, these findings reveal that cognate peptide binding suppresses intermediate timescale conformational exchange processes across the active site of RASProtease(II) while inducing distinct dynamic effects at distal regions, whereas non-cognate binding increases conformational exchange both locally and distally. These contrasting dynamic responses provide a fundamental basis for understanding differences in catalytic efficiency as discussed further below.

## Discussion

The general mechanisms responsible for distinguishing the activities of cognate and off-target non-cognate substrate sequences in protease-catalyzed reactions are still not well understood. Indeed, the distinction between cognate and non-cognate sequences can be quite subtle, where even a relatively small, single amino acid, change can lead to large differences in catalytic activity. A striking example of this subtlety is the designed RASProtease(II), which targets the active GTP-bound state of RAS at the switch II cognate sequence, QEEYSAM, with a ∼30-fold higher activity than the non-cognate sequence, QEEISAM.

Despite the substantial difference in catalytic efficiency between the cognate and non-cognate substrates, RASProtease(II) retains a remarkably consistent structure upon substrate binding. Structural alignment of both molecules in the asymmetric unit (536 residues total) to the apo structure shows that the protease exhibits minimal global structural differences for both the cognate QEEYSAM and non-cognate QEEISAM complexes. Only 3 residues in the cognate complex and 1 residue in the non-cognate complex deviate by more than 0.90 Å. These values correspond to well under 1% positional change in each comparison. Similarly, direct alignment of the asymmetric units of the QEEYSAM and QEEISAM complexes reveals only 1 residue with positional deviation at the same RMSD cutoff, further emphasizing the high structural conservation between the peptide-bound forms. Beyond global structural alignment, analysis of electron density reveals localized conformational differences between the two complexes. These subtle differences in alternate conformations are limited to roughly 9% of residues, including S128, which lies at the protease-peptide interface and undergoes changes in conformational dynamics. Taken together, the minimal structural deviations and limited changes to alternate conformations indicate that the overall fold of RASProtease(II) is highly conserved upon substrate binding, and this would suggest that the differences in catalytic efficiency arise primarily from local interactions at the substrate interface, particularly at the P4 residue, which is the farthest peptide-docking position relative to the catalytic triad. The larger tyrosine in the cognate sequence packs more favorably than the smaller isoleucine in the non-cognate, likely increasing local stability of the S4 pocket.

However, it was not clear from the crystal structures how differences at the P4 position led to large variations in activity at the proteolytic cleavage site, particularly as both the cognate and non-cognate complexes form the acyl bond between S221 and the peptide. Thus, the contrast in activity appears to have more complex origins than simple differences in local interactions in the vicinity of the S4 pocket. NMR-based binding experiments provided a clue by revealing that the cognate and non-cognate peptides induce distinct long-range CSP and peak intensity effects in RASProtease(II). Based on these observations, and insights into the role of dynamic motions in enzyme catalysis (41), we hypothesized that the differences in catalytic activity are due to substrate-induced effects on protease conformational dynamics. Employing both X-ray crystallographic anisotropic displacement parameter analysis and solution NMR spectroscopy, our results support this postulate and demonstrate significant changes in RASProtease(II) dynamics between the cognate and non-cognate forms.

Crystallographic anisotropy analysis of the peptides themselves showed that the cognate QEEYSAM peptide undergoes limited motion in the binding pocket, due to multiple stabilizing interactions with the protease, whereas the non-cognate QEEISAM peptide remains more conformationally dynamic to compensate for fewer favorable contacts. These differences are mirrored in the protease, with the cognate complex showing reduced anisotropic motion in several helical regions and the non-cognate complex retaining greater flexibility (**Figure 3)**. Notably, even subtle changes induced by the non-cognate peptide propagate to residues distal from the active site, highlighting its broader impact on protease dynamics. These differential effects underscore how substrate identity modulates enzyme dynamics and are further supported by NMR experiments.

NMR analysis of RASProtease(II) showed limited differences in fast timescale (ps–ns) dynamics between the cognate and non-cognate complexes. However, significant changes were observed at slower timescales (µs–ms) through ^15^N CPMG RD and CEST experiments. The apo protease displays extensive conformational dynamics, which are quenched around the active site in the cognate QEEYSAM complex. Most backbone amides of residues lining the S1–S4 pockets show either decreased conformational exchange (Δ*R*_ex_ < 0) values or become detectable in NMR spectra of the cognate complex. This kind of local stabilization and quenching of dynamics at the protease-peptide interface has also been observed in other proteases, such as thrombin binding to the inhibitor PPACK (12). In addition, binding of the cognate peptide to RASProtease(II) enhances conformational exchange in many distal residues organized into three clusters that each span from the active site to multiple exterior regions of the protease (**Figure 7A)**. Cluster (i) extends from N64 near the S2 pocket to C87 in a loop ∼25 Å away. Cluster (ii) incorporates A153 near the S1 site and extends to S159 (loop) and K136 (helix), both more than 15 Å away from S1. Cluster (iii) starts at T73 near the S1 pocket and primarily involves residues from several helices, including Q19, K237, and V246, each located more than 25 Å from S1. Thus, while binding of QEEYSAM stabilizes residues near the active site, it also induces destabilization of residues around the S1 and S2 pockets that propagates to more distal regions of the protease. These changes redistribute dynamics from the active site to more remote structural elements, thereby engaging a broader allosteric network and likely compensating for the entropic cost of active site stabilization.

In contrast, interaction with the non-cognate QEEISAM peptide leads to a completely different dynamic profile for RASProtease(II). The stabilizing effects of the non-cognate peptide are much smaller, with only a few residues in the active site exhibiting reduced conformational exchange. Most residues in this region, however, either retain levels of exchange similar to the apo state or become further destabilized (**Figure 7B)**. The most significant destabilized network, cluster (iv), begins at S101 near the S3 site and extends along the substrate-binding groove through S2 and S1, involving proximal residues D99 and S218, respectively. It ultimately perturbs G219 in the S1 site, located at the N-terminus of the central helix (220–237), where exchange-broadening is so severe that its signal becomes undetectable by NMR. Smaller networks, clusters (v) and (vi), are analogous to the cognate clusters (ii) and (iii), respectively, and highlight destabilization adjacent to the central helix that propagates from the S1 pocket to more distal regions of the protease. Beyond the destabilized clusters, certain residues illustrate pronounced substrate-dependent differences in conformational dynamics. For example, S128 in the S4 pocket has dampened motion in the cognate QEEYSAM complex, but remains exchange-broadened in both the apo and non-cognate states, reflecting substantial conformational heterogeneity. This dynamic behavior is corroborated by the crystal structures, where S128 adopts alternate conformations clearly visible in the electron density maps.

Together, these observations indicate that the mutation from a tyrosine to an isoleucine in the substrate peptide induces destabilizing effects in the S4 pocket of RASProtease(II) that also propagate allosterically throughout the protease S1–S4 pockets to the catalytic site. Conformational destabilization in these protease regions—particularly at the N-terminus of the central helix, where the catalytic S221 residue forms the acyl intermediate with the substrate—is likely to hinder the acylation step, providing a plausible explanation for why cleavage of the non-cognate QEEISAM peptide is ∼30-fold slower than that of the cognate substrate. The NMR RD findings are further supported by the X-ray anisotropy and NMR CEST analyses, both of which indicate that the S1–S4 region adopts a predominantly homogeneous conformation in the cognate QEEYSAM complex, but exhibits pronounced conformational heterogeneity in the non-cognate QEEISAM complex. Thus, while the cognate peptide quenches conformational dynamics in the peptide-docking subsites and active site to facilitate optimal proteolytic activity, the non-cognate peptide not only exerts a diminished stabilizing effect, but also increases conformational dynamics in these regions.

In conclusion, we provide here a detailed characterization of the kinetic, structural, and dynamic features for cognate and non-cognate complexes with RASProtease(II), a protease engineered to target RAS, a key signaling protein implicated in many cancers. Our results indicate that dynamic allostery plays a major role in modulating RASProtease(II) reactivity, revealing important insights into the molecular basis for differentiating between cognate and non-cognate substrates. Although these conclusions reflect the product-bound state prior to its release, examination of additional intermediates along the enzymatic reaction coordinate in future studies will provide a broader basis for comparison with the behavior described here. Moreover, because RASProtease(II) is derived from the prototypical serine protease, subtilisin, it is likely that the overall mechanism employed for selecting cognate over off-target non-cognate sequences presented here will be generalizable to many other proteases. In particular, from a protein design perspective, it may be possible to further modulate RASProtease selectivity by targeting dynamic allosteric networks, including residues distal to the active site. Such strategies, if successful, would expand the target sequence space available for mutation and provide a major advance in the rational design of protease specificity. A further indication of the potential of this approach is exhibited by the thrombin-thrombomodulin complex, in which thrombomodulin binds at a site more than 25 Å away from the S1 pocket of thrombin, yet allosterically rigidifies that pocket to promote protein C cleavage (13). Notably, these experimental findings are expected to support the validation of computational approaches for simulating the differential physicochemical responses of proteases to cognate versus non-cognate substrates (1, 42), thereby accelerating the field of enzyme design.

## Materials and Methods

### Protein Expression and Purification

The prodomain-RASProtease(II) construct was cloned into the pJ1 expression vector (43), transformed into *E. coli* BL21(DE3) cells (New England Biolabs), and grown in either LB or M9 minimal media at 37°C until an OD_600_ of 0.6–0.8 was reached.

Standard procedures were used for labeling of RASProtease(II) with stable isotopes (^2^H, ^13^C, ^15^N) in minimal media. Cells were first adapted to growth in 30% ^2^H_2_O, gradually incrementing to 100% ^2^H_2_O. Protein expression was induced with 1 mM IPTG, typically taking 3–4 hours in LB media, 6–8 hours in M9 media (^15^N labeling), and 24–48 hours in highly deuterated media, followed by overnight growth at 25°C.

Upon collection of the cells, pellets were resuspended in buffer (100 mM potassium phosphate, 150 mM sodium chloride, pH 7.0) supplemented with EDTA-free protease inhibitor tablets (Thermo Fisher Scientific) and lysed using sonication. The lysate was cleared by centrifugation and the supernatant fraction was incubated with 10 mM imidazole at 25°C for 1 hour to cleave the prodomain. The sample was then loaded onto a 1 mL N-hydroxysuccinimide column (Cytiva) derivatized to the N-terminus of the cognate QEEYSAM peptide (AnaSpec). The column was washed as follows: 5 column volumes (CV) of running buffer (100 mM potassium phosphate, pH 7.0), 20 CV of wash buffer (100 mM potassium phosphate, 100 mM sodium chloride, pH 7.2), 5 CV of running buffer, and finally, 10 CV of elution buffer (10 mM ammonium hydroxide solution). Fractions were neutralized using 1 M potassium phosphate buffer pH 7.0 to a final concentration of 100 mM and analyzed by SDS-PAGE. Pure fractions were combined and concentrated for subsequent experiments.

For the triple-labeled RASProtease(II) sample, an additional refolding step was employed to back-exchange buried backbone amides. RASProtease(II) was treated with 6 M guanidinium hydrochloride at room temperature for 2 hours. The protein solution was then added dropwise into a stirred solution of 200 mM potassium phosphate, pH 7.5 (∼17 mL), and allowed to refold at room temperature for 1 hour.

RASProtease(II) samples were buffer exchanged into 100 mM potassium phosphate, 0.5 mM EDTA, pH 7.0, and concentrated to 300 µM for NMR analysis.

### Crystallization and Data Collection

RASProtease(II) was buffer exchanged into 100 mM potassium phosphate, pH 7.0 and concentrated to 150 µM for crystallization screening. The best crystals were obtained in a condition containing 200 mM sodium formate and 20% PEG 3350.

The QEEYSAM and QEEISAM peptides were purchased from AnaSpec and resuspended at 50 mM in DMSO. The QEEYSAM-bound RASProtease(II) and QEEISAM-bound RASProtease(II) complexes were co-crystallized by incubating the protease with the peptide at a 1:1 molar ratio for 15 minutes at room temperature prior to setting up drops. Crystals of both complexes were obtained in a condition containing 50 mM zinc acetate and 15% PEG 3350.

All crystals were grown at 22°C, appearing overnight in a rod-like morphology and reaching a maximum size within 3–7 days. Crystals were transferred to the crystallization condition supplemented with 20–25% glycerol and cryo-cooled in liquid nitrogen prior to data collection.

Diffraction data for the apo RASProtease(II) and the QEEYSAM-bound RASProtease(II) complex were collected at the Southeast Regional Collaborative Access Team (SER-CAT) 22-ID-D beamline at the Advanced Photon Source, Argonne National Laboratory. Diffraction data for the QEEISAM-bound RASProtease(II) complex were collected at the Frontier Microfocusing Macromolecular Crystallography (FMX) 17-ID-2 beamline at the National Synchrotron Light Source II, Brookhaven National Laboratory (44).

### Structure Determination, Model Building, and Refinement

Diffraction data were indexed, integrated, and scaled using HKL2000 (45), and subsequently merged in Aimless (46, 47). Initial phases were determined by molecular replacement using the progenitor RASProtease model (PDB ID: 6U9L) (4) as the search model through Phaser (48) in the CCP4 suite (49). Prior to molecular replacement, all solvent molecules and ions were removed from the search model. The structure was initially refined using Refmac (50, 51) in the CCP4 suite (49). Subsequent refinement was conducted in Phenix (52) and iterative cycles of model building were performed in Coot (53, 54). Final refinement statistics are shown in Table S1. The refined structures were deposited in the PDB under the following accession codes: 9ZIO, 9ZIP, and 9ZIQ.

### Crystallographic Anisotropy Analysis

The final crystallographic models were analyzed in Anisoanl from the CCP4 suite (47) to obtain the anisotropic values for each residue. Values near 1.0 indicate more isotropic (uniform) motion, whereas values closer to 0.0 reflect high anisotropy. Z-scores were calculated for the anisotropy values of the apo RASProtease(II) structure and the top 20% most anisotropic residues were identified based on this distribution. These residues were then used as reference points to compare the anisotropy in the QEEYSAM-bound and QEEISAM-bound complexes. Significant residues were identified and mapped onto the corresponding crystal structures for visual comparison using PyMOL (55) and UCSF Chimera (56).

### Stopped-flow Kinetics

Kinetic measurements were conducted using the Applied Photophysics SX20 stopped flow spectrometer and the Applied Photophysics Pro-Data SX software (version 2.5.1852.0). Fluorescence was detected by a photomultiplier tube mounted at a 90° angle to incident light. Experiments were conducted at an excitation wavelength of 360 nm with a 400 nm energy filter for detection of the 430 nm emission peak, based on the intrinsic properties of the amino-4-methylcoumarin (AMC) fluorophore.

The QEEYSAM-AMC and QEEISAM-AMC substrates were purchased from AnaSpec and resuspended at 25 mM in DMSO. Immediately before the experiment, the peptides were diluted to desired concentrations in 100 mM potassium phosphate pH 7.0 buffer containing 10 mM imidazole. RASProtease(II) samples were prepared in the same buffer. The protease and peptide samples were prepared and loaded separately into 3 mL syringes and rapidly mixed at a 1:1 volumetric ratio by the stopped-flow instrument at room temperature to make the final solutions at desired concentrations (100 nM for the substrate and 4 µM for the protease). For each scan, approximately 120 µL was injected through the optical cell and measured over 1800 seconds, with each scan generating 10,000 data points.

Experiments were conducted under single-turnover conditions, with enzyme concentration in excess over substrate concentration ([E] >> [S]). The observed fluorescence signal, which is directly proportional to the fraction of product formed, was normalized such that the maximum fluorescence corresponded to complete product formation. Background fluorescence from control samples containing substrate and buffer (without protease) was subtracted prior to normalization. For each condition, three replicate measurements were collected. Fluorescence progress curves (Relative Fluorescence vs. Time) were fitted by non-linear regression to Equation (1) using MATLAB (57). The resulting *k*_obs_ values were averaged to report the mean ± 1 standard deviation.

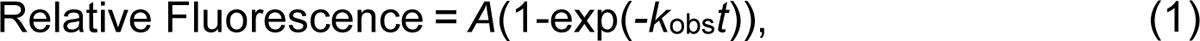

where *A* is the amplitude, *k*_obs_ is the rate constant, and *t* is the reaction time.

### Nuclear Magnetic Resonance (NMR) Spectroscopy

NMR data were collected at 37°C on Bruker Avance III 600 MHz and 900 MHz spectrometers fitted with a Z-gradient triple resonance [^1^H/^13^C/^15^N]-cryoprobe.

Backbone resonances were assigned for the apo and QEEYSAM-bound RASProtease(II) using TROSY-based three-dimensional HNCACB, HN(CO)CACB, HNCO, and HN(CA)CO experiments with 25% non-uniform sampling (58). Main chain amide assignments for the QEEISAM-bound RASProtease(II) were obtained by comparison with the apo and QEEYSAM-bound two-dimensional ^1^H-^15^N HSQC spectra.

Spectra were processed using NMRPipe (59) and analyzed with Sparky (60). NMR spectra of stoichiometric 1:1 complexes between RASProtease(II) and QEEYSAM or QEEISAM peptides were obtained using the same conditions as for the apo state.

Peptides were added to RASProtease(II) from a 50 mM stock solution in DMSO. Backbone amide chemical shift perturbations upon peptide additions were calculated by employing Equation (2).

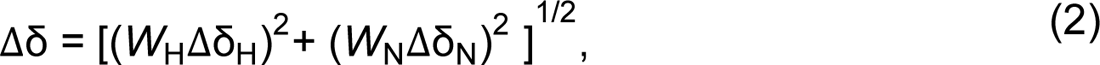

where Δδ_H_ and Δδ_N_ are the ^1^H and ^15^N chemical shift differences for a specific signal, respectively, and *W*_H_=1 and *W*_N_=0.2 are weighting factors.

^15^N-Spin-lattice (*R*_1_), ^15^N-spin-spin (*R*_2_), and steady state {^1^H}-^15^N heteronuclear NOE experiments were recorded on RASProtease(II) samples in the unbound and peptide-bound states at 37°C on a 600 MHz spectrometer. Heteronuclear NOE data were collected with a relaxation delay of 3 s in interleaved mode. Background noise level was used to estimate errors in heteronuclear NOE values. ^15^N-*R*_1_ and ^15^N-*R*_2_ measurements were carried out using a relaxation delay of 3.5 s. For *R*_1_ experiments, the following variable delays were used: 50, 100, 200, 300, 400, 600, 800, 1000, 1400, and 2000 ms. The variable delays for *R*_2_ experiments were: 5, 10, 15, 30, 40, 60, 70, 100, 140, and 200 ms. Both *R*_1_ and *R*_2_ values were obtained by fitting an exponential decay with error estimates utilizing Sparky (60).

TROSY-based ^15^N constant-time CPMG relaxation dispersion experiments were collected with temperature compensation in pseudo-3D mode (61, 62). CPMG field strengths of 0–1000 Hz were used with a 60 ms constant-time relaxation period, 2 s relaxation delay, 128 *t*_1_ increments, and 128 transients per FID. Spectra were processed with NMRPipe (59) and Sparky peak lists were imported into NESSY (63) for automated fitting of estimated *R*_ex_ values and analysis. Error bars for the cognate QEEYSAM and non-cognate QEEISAM complexes were obtained in NESSY from 3 duplicate RD experiments. Error bars for the apo RD data were estimated by comparing noise levels with those of the peptide-bound datasets.

TROSY-based ^15^N-CEST experiments (64) were acquired at 600 MHz in a pseudo-3D mode employing a *B*_1_ field strength of 25 Hz and a relaxation delay *T*_relax_ of 400 ms. ^15^N-Offsets were between 103–134 ppm in 0.5 ppm increments. Spectra were collected with 44–52 scans per FID and a recycle delay of 1.0 s. Acquired pseudo-3D data were split into 64 x 2D experiments and processed with NMRPipe (59). CEST curves were obtained by plotting the peak intensity ratios (*I*/*I*_0_) versus the ^15^N-offset frequency for the backbone amide peak of each residue, where *I* is the intensity at each offset and *I*_0_ is the reference intensity at 103 ppm. Differences in the CEST profile for a given residue (Δω, Hz) were calculated relative to the apo state for both the cognate and non-cognate forms, using the width at half-height of the major *I*/*I*_0_ dip in each profile.

## Author Contributions

**Betty Chu:** Investigation; methodology; formal analysis; visualization; writing – original draft; writing – review and editing; data curation. **Yanan He:** Investigation; formal analysis. **Yihong Chen:** Investigation. **Eric A. Toth:** Conceptualization; supervision; funding acquisition; writing – original draft; writing – review and editing; project administration; validation. **John Orban:** Conceptualization; supervision; funding acquisition; writing – original draft; writing – review and editing; formal analysis; visualization; project administration; validation.

## Acknowledgements

This research was financially supported by the National Institute of General Medical Sciences, Grants 5R01GM141290-03 and 3R01GM141290-03S1, awarded to J.O and E.A.T. We acknowledge Dr. Philip N. Bryan, Potomac Affinity Proteins, MD, USA, for providing the sequence information and initial supply of RASProtease(II). We thank Dr. Jaekyun Jeon, Institute for Bioscience and Biotechnology Research (IBBR), University of Maryland, for providing access to the stopped-flow instrument used in the kinetics experiments. We also thank Dr. Alexander C. Drohat, University of Maryland School of Medicine, for helpful discussions that contributed to the interpretation of the kinetics data. Crystallographic data were collected at the Southeast Regional Collaborative Access Team (SER-CAT) 22-ID-D beamline at the Advanced Photon Source (APS), Argonne National Laboratory (ANL) and at the FMX17-ID-2 beamline at the National Synchrotron Light Source II (NSLS-II), Brookhaven National Laboratory (BNL). SER-CAT is supported by its member institutions and equipment grants (S10_RR25528, S10_RR028976 and S10_OD027000) from the National Institutes of Health (NIH). As part of the APS, a national user facility at ANL, work conducted at SER-CAT is supported by the U.S. Department of Energy (DOE) Office of Science under Contract No. DE-AC02-06CH11357. The FMX beamline at NSLS-II is primarily supported by the NIH National Institute of General Medical Sciences (NIGMS) through a Center Core P30 Grant (P30GM133893) and by the DOE Office of Biological and Environmental Research (KP1605010). As part of NSLS-II, a national user facility at BNL, work conducted at FMX is supported by the U.S. DOE Office of Science under Contract No. DE-SC0012704. The NMR facility at IBBR is supported by the University of Maryland, the National Institute of Standards and Technology, and a grant from the W. M. Keck Foundation.

## Data Availability Statement

The data that support the findings of this study are available in the supplementary material of this article. Crystallographic structures are available from the RCSB Protein Data Bank under the accession codes 9ZIO, 9ZIP, and 9ZIQ. NMR spectra are available from the Biological Magnetic Resonance Bank under the accession codes 53227, 53228, and 53229.

## References

1. Dyer RP & Weiss GA (2022) Making the cut with protease engineering. Cell Chem Biol 29(2):177–190.

2. Packer MS, Rees HA, & Liu DR (2017) Phage-assisted continuous evolution of proteases with altered substrate specificity. Nat Commun 8(1):956.

3. Craik CS, Page MJ, & Madison EL (2011) Proteases as therapeutics. Biochem J 435(1):1–16.

4. Chen Y, et al. (2021) Engineering subtilisin proteases that specifically degrade active RAS. Commun Biol 4(1):299.

5. Gron H, Meldal M, & Breddam K (1992) Extensive comparison of the substrate preferences of two subtilisins as determined with peptide substrates which are based on the principle of intramolecular quenching. Biochemistry 31(26):6011–6018.

6. Rockwell NC, Krysan DJ, Komiyama T, & Fuller RS (2002) Precursor processing by kex2/furin proteases. Chem Rev 102(12):4525–4548.

7. Cui Q & Karplus M (2008) Allostery and cooperativity revisited. Protein Sci 17(8):1295–1307.

8. Gunasekaran K, Ma B, & Nussinov R (2004) Is allostery an intrinsic property of all dynamic proteins? Proteins 57(3):433–443.

9. Hauske P, Ottmann C, Meltzer M, Ehrmann M, & Kaiser M (2008) Allosteric regulation of proteases. Chembiochem 9(18):2920–2928.

10. Swain JF & Gierasch LM (2006) The changing landscape of protein allostery. Curr Opin Struct Biol 16(1):102–108.

11. Cooper A & Dryden DT (1984) Allostery without conformational change. A plausible model. Eur Biophys J 11(2):103–109.

12. Handley LD, et al. (2017) NMR reveals a dynamic allosteric pathway in thrombin. Sci Rep 7:39575.

13. Peacock RB, McGrann T, Tonelli M, & Komives EA (2021) Serine protease dynamics revealed by NMR analysis of the thrombin-thrombomodulin complex. Sci Rep 11(1):9354.

14. Valles GJ, et al. (2025) Activation dynamics of ubiquitin-specific protease 7. Proc Natl Acad Sci U S A 122(21):e2426632122.

15. Guo J & Zhou HX (2016) Protein Allostery and Conformational Dynamics. Chem Rev 116(11):6503–6515.

16. Burns MC, et al. (2014) Approach for targeting Ras with small molecules that activate SOS-mediated nucleotide exchange. Proc Natl Acad Sci U S A 111(9):3401–3406.

17. Canon J, et al. (2019) The clinical KRAS(G12C) inhibitor AMG 510 drives anti-tumour immunity. Nature 575(7781):217–223.

18. Hallin J, et al. (2020) The KRAS(G12C) Inhibitor MRTX849 Provides Insight toward Therapeutic Susceptibility of KRAS-Mutant Cancers in Mouse Models and Patients. Cancer Discov 10(1):54–71.

19. Janes MR, et al. (2018) Targeting KRAS Mutant Cancers with a Covalent G12C-Specific Inhibitor. Cell 172(3):578–589 e517.

20. Ostrem JM, Peters U, Sos ML, Wells JA, & Shokat KM (2013) K-Ras(G12C) inhibitors allosterically control GTP affinity and effector interactions. Nature 503(7477):548–551.

21. Ostrem JM & Shokat KM (2016) Direct small-molecule inhibitors of KRAS: from structural insights to mechanism-based design. Nat Rev Drug Discov 15(11):771–785.

22. Patricelli MP, et al. (2016) Selective Inhibition of Oncogenic KRAS Output with Small Molecules Targeting the Inactive State. Cancer Discov 6(3):316–329.

23. Skoulidis F, et al. (2021) Sotorasib for Lung Cancers with KRAS p.G12C Mutation. N Engl J Med 384(25):2371–2381.

24. Hallin J, et al. (2022) Anti-tumor efficacy of a potent and selective non-covalent KRAS(G12D) inhibitor. Nat Med 28(10):2171–2182.

25. Zhou C, et al. (2024) Anti-tumor efficacy of HRS-4642 and its potential combination with proteasome inhibition in KRAS G12D-mutant cancer. Cancer Cell 42(7):1286–1300 e1288.

26. Holderfield M, et al. (2024) Concurrent inhibition of oncogenic and wild-type RAS-GTP for cancer therapy. Nature 629(8013):919–926.

27. Jiang J, et al. (2024) Translational and Therapeutic Evaluation of RAS-GTP Inhibition by RMC-6236 in RAS-Driven Cancers. Cancer Discov 14(6):994–1017.

28. Kessler D, et al. (2019) Drugging an undruggable pocket on KRAS. Proc Natl Acad Sci U S A 116(32):15823–15829.

29. Kim D, et al. (2023) Pan-KRAS inhibitor disables oncogenic signalling and tumour growth. Nature 619(7968):160–166.

30. Wasko UN, et al. (2024) Tumour-selective activity of RAS-GTP inhibition in pancreatic cancer. Nature 629(8013):927–936.

31. Ma Y, et al. (2013) Targeted degradation of KRAS by an engineered ubiquitin ligase suppresses pancreatic cancer cell growth in vitro and in vivo. Mol Cancer Ther 12(3):286–294.

32. Nagashima T, et al. (2022) ASP3082, a First-in-class novel KRAS G12D degrader, exhibits remarkable anti-tumor activity in KRAS G12D mutated cancer models. European Journal of Cancer 174:S30.

33. Biancucci M, et al. (2018) The bacterial Ras/Rap1 site-specific endopeptidase RRSP cleaves Ras through an atypical mechanism to disrupt Ras-ERK signaling. Sci Signal 11(550).

34. Biancucci M, et al. (2017) Substrate Recognition of MARTX Ras/Rap1-Specific Endopeptidase. Biochemistry 56(21):2747–2757.

35. Biancucci M & Satchell KJ (2015) A bacterial toxin that cleaves Ras oncoprotein. Oncotarget 6(22):18742–18743.

36. Escher TE, et al. (2025) Expression of a RAS Degrader via Synthetic Nanocarrier-Mediated mRNA Delivery Reduces Pancreatic Tumors. ACS Appl Bio Mater 8(6):4805-4814.

37. Stubbs CK, Biancucci M, Vidimar V, & Satchell KJF (2021) RAS specific protease induces irreversible growth arrest via p27 in several KRAS mutant colorectal cancer cell lines. Sci Rep 11(1):17925.

38. Vidimar V, et al. (2020) An engineered chimeric toxin that cleaves activated mutant and wild-type RAS inhibits tumor growth. Proc Natl Acad Sci U S A 117(29):16938–16948.

39. Vidimar V, et al. (2022) Proteolytic pan-RAS Cleavage Leads to Tumor Regression in Patient-derived Pancreatic Cancer Xenografts. Mol Cancer Ther 21(5):810–820.

40. Schechter I & Berger A (1967) On the size of the active site in proteases. I. Papain. Biochem Biophys Res Commun 27(2):157–162.

41. Deshmukh L, Tugarinov V, Louis JM, & Clore GM (2017) Binding kinetics and substrate selectivity in HIV-1 protease-Gag interactions probed at atomic resolution by chemical exchange NMR. Proc Natl Acad Sci U S A 114(46):E9855–E9862.

42. Lu C, et al. (2023) Prediction and design of protease enzyme specificity using a structure-aware graph convolutional network. Proc Natl Acad Sci U S A 120(39):e2303590120.

43. Ruan B, Hoskins J, & Bryan PN (1999) Rapid folding of calcium-free subtilisin by a stabilized pro-domain mutant. Biochemistry 38(26):8562–8571.

44. Schneider DK, et al. (2021) FMX - the Frontier Microfocusing Macromolecular Crystallography Beamline at the National Synchrotron Light Source II. J Synchrotron Radiat 28(Pt 2):650–665.

45. Otwinowski Z & Minor W (1997) Processing of X-ray diffraction data collected in oscillation mode. Methods Enzymol 276:307–326.

46. Evans PR & Murshudov GN (2013) How good are my data and what is the resolution? Acta Crystallogr D Biol Crystallogr 69(Pt 7):1204–1214.

47. Winn MD, et al. (2011) Overview of the CCP4 suite and current developments. Acta Crystallogr D Biol Crystallogr 67(Pt 4):235–242.

48. McCoy AJ, et al. (2007) Phaser crystallographic software. J Appl Crystallogr 40(Pt 4):658–674.

49. Potterton L, et al. (2018) CCP4i2: the new graphical user interface to the CCP4 program suite. Acta Crystallogr D Struct Biol 74(Pt 2):68–84.

50. Murshudov GN, et al. (2011) REFMAC5 for the refinement of macromolecular crystal structures. Acta Crystallogr D Biol Crystallogr 67(Pt 4):355–367.

51. Murshudov GN, Vagin AA, & Dodson EJ (1997) Refinement of macromolecular structures by the maximum-likelihood method. Acta Crystallogr D Biol Crystallogr 53(Pt 3):240–255.

52. Liebschner D, et al. (2019) Macromolecular structure determination using X-rays, neutrons and electrons: recent developments in Phenix. Acta Crystallogr D Struct Biol 75(Pt 10):861–877.

53. Emsley P & Cowtan K (2004) Coot: model-building tools for molecular graphics. Acta Crystallogr D Biol Crystallogr 60(Pt 12 Pt 1):2126–2132.

54. Emsley P, Lohkamp B, Scott WG, & Cowtan K (2010) Features and development of Coot. Acta Crystallogr D Biol Crystallogr 66(Pt 4):486–501.

55. Schrodinger, LLC (2015) The PyMOL Molecular Graphics System, Version 3.1.3.

56. Pettersen EF, et al. (2004) UCSF Chimera--a visualization system for exploratory research and analysis. J Comput Chem 25(13):1605–1612.

57. The MathWorks Inc. (2024) MATLAB Version 24.2.0 (R2024b) (The MathWorks Inc., Natick, Massachusetts).

58. Hyberts SG, Arthanari H, & Wagner G (2012) Applications of non-uniform sampling and processing. Top Curr Chem 316:125–148.

59. Delaglio F, et al. (1995) NMRPipe: a multidimensional spectral processing system based on UNIX pipes. J Biomol NMR 6(3):277–293.

60. Goddard T & Kneller D (2021) SPARKY 3 (University of California, San Francisco, 2008).

61. Hansen DF, Vallurupalli P, & Kay LE (2008) An improved 15N relaxation dispersion experiment for the measurement of millisecond time-scale dynamics in proteins. J Phys Chem B 112(19):5898–5904.

62. Jiang B, Yu B, Zhang X, Liu M, & Yang D (2015) A (15)N CPMG relaxation dispersion experiment more resistant to resonance offset and pulse imperfection. J Magn Reson 257:1–7.

63. Bieri M & Gooley PR (2011) Automated NMR relaxation dispersion data analysis using NESSY. BMC Bioinformatics 12:421.

64. Long D, Bouvignies G, & Kay LE (2014) Measuring hydrogen exchange rates in invisible protein excited states. Proc Natl Acad Sci U S A 111(24):8820–8825.

