## Supplementary Information for "Substrate specificity in a designed RAS-targeting protease is coupled to active site and distal motions"

### Supplementary Table

#### Table S1. Data Collection and Refinement Statistics.*

| | **Apo RASProtease(II)** | **QEEYSAM-bound RASProtease(II)** | **QEEISAM-bound RASProtease(II)** |
|---|---|---|---|
| **PDB ID** | 9ZIO | 9ZIP | 9ZIQ |
| **Data Collection** | | | |
| Space Group | $P12_11$ | $P12_11$ | $P12_11$ |
| Cell Dimensions | | | |
| a, b, c (Å) | 49.13, 60.58, 70.16 | 43.98, 57.91, 82.49 | 44.24, 58.24, 82.60 |
| α, β, γ (°) | 90.00, 93.75, 90.00 | 90.00, 98.06, 90.00 | 90.00, 97.73, 90.00 |
| Resolution (Å) | 45.81–1.49 (1.52–1.49) | 47.24–1.24 (1.26–1.24) | 58.24–1.20 (1.22–1.20) |
| I/σ(I) | 15.8 (2.4) | 18.0 (4.4) | 7.7 (1.6) |
| Completeness (%) | 97.9 (91.8) | 94.0 (86.8) | 98.3 (94.8) |
| Multiplicity | 6.5 (6.2) | 6.0 (5.8) | 5.5 (4.7) |
| **Refinement** | | | |
| Resolution (Å) | 41.45–1.49 | 47.24–1.24 | 40.92–1.20 |
| $R_{work}/R_{free}$ (%) | 12.21/15.85 | 11.27/13.97 | 13.54/15.96 |
| No. atoms | 4470 | 4786 | 4603 |
| Protein | 3891 | 4074 | 3934 |
| Ligand/Ion | 103 | 81 | 128 |
| Water | 476 | 631 | 541 |
| Anisotropy | 0.123 | 0.263 | 0.147 |
| Average B-factors ($Å^2$) | 18.0 | 13.0 | 16.0 |
| RMS deviations | | | |
| Bonds (Å) | 0.008 | 0.013 | 0.013 |
| Angles (°) | 1.058 | 1.622 | 1.373 |

*Values in parentheses correspond to the highest resolution shell.

### Supplementary Figures

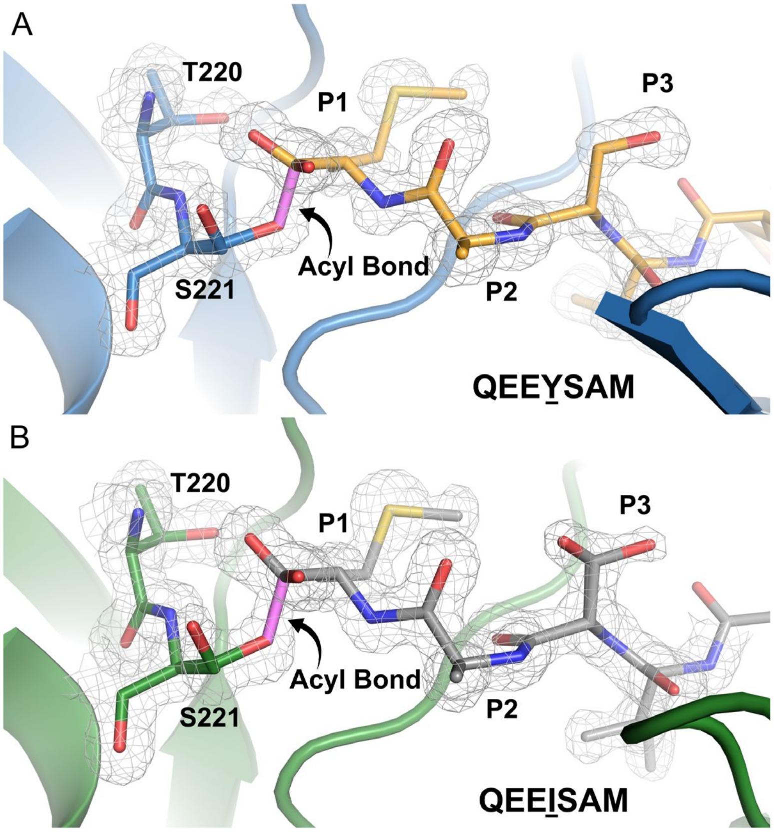


**Figure S1. Formation of the acyl-enzyme intermediate in peptide-bound RASProtease(II) complexes.** Residues T220 and S221 of RASProtease(II) and residues P1 through P3 of the peptide are shown with $2mF_o-DF_c$ electron density contoured at 1 σ. **(A)** The cognate QEEYSAM complex. The protease and peptide atoms are colored blue and orange, respectively. **(B)** The non-cognate QEEISAM complex. The protease and peptide atoms are colored green and light gray, respectively. The acyl bond between the serine and methionine is highlighted in violet in both complexes.

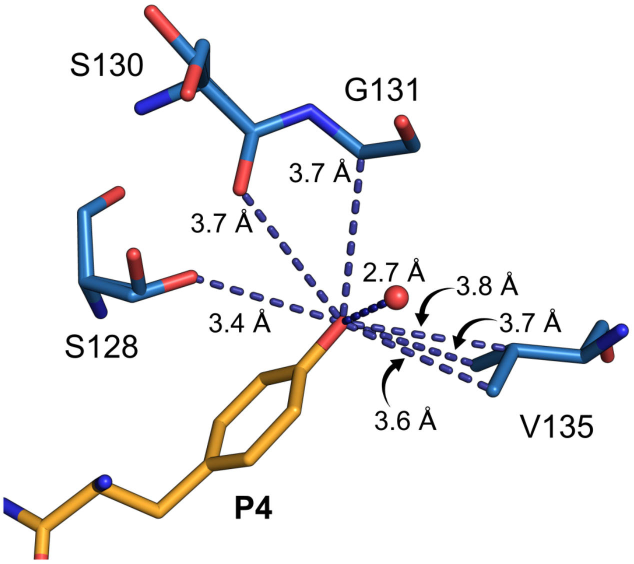


**Figure S2. Contacts between the P4 hydroxyl group and RASProtease(II) in the cognate complex.** The tyrosine OH atom of the cognate QEEYSAM peptide coordinates a water molecule (colored red) at 2.7 Å and hydrogen bonds with residue S128 in RASProtease(II). The hydroxyl group forms additional van der Waals interactions with residues S130, G131, and V135 at 3.4–3.8 Å distances.

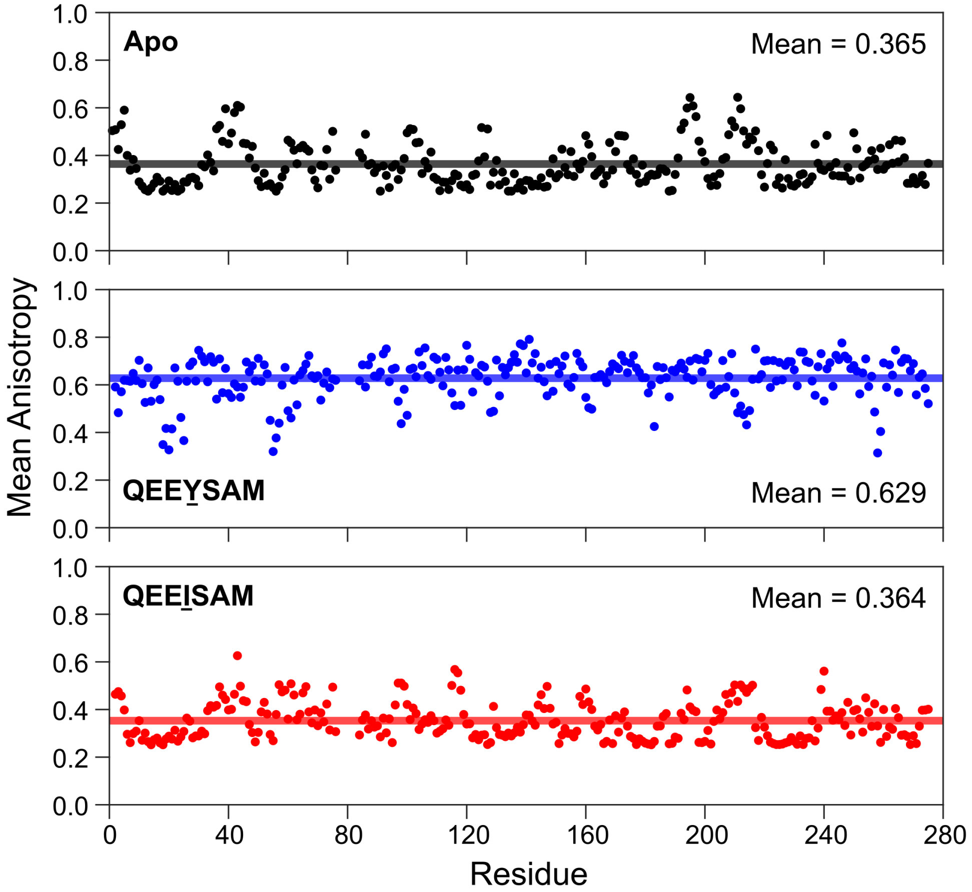


**Figure S3. Anisotropy of the main chain protease atoms of the apo, QEEYSAM-bound, and QEEISAM-bound RASProtease(II) structures.** Anisotropy values are plotted for each residue in the apo form (top), cognate QEEYSAM complex (center), and non-cognate QEEISAM complex (bottom). The mean anisotropy value for each structure is indicated by a horizontal line and labeled in each graph. Values closer to 1.0 reflect more isotropic motion, whereas smaller values indicate greater anisotropy.

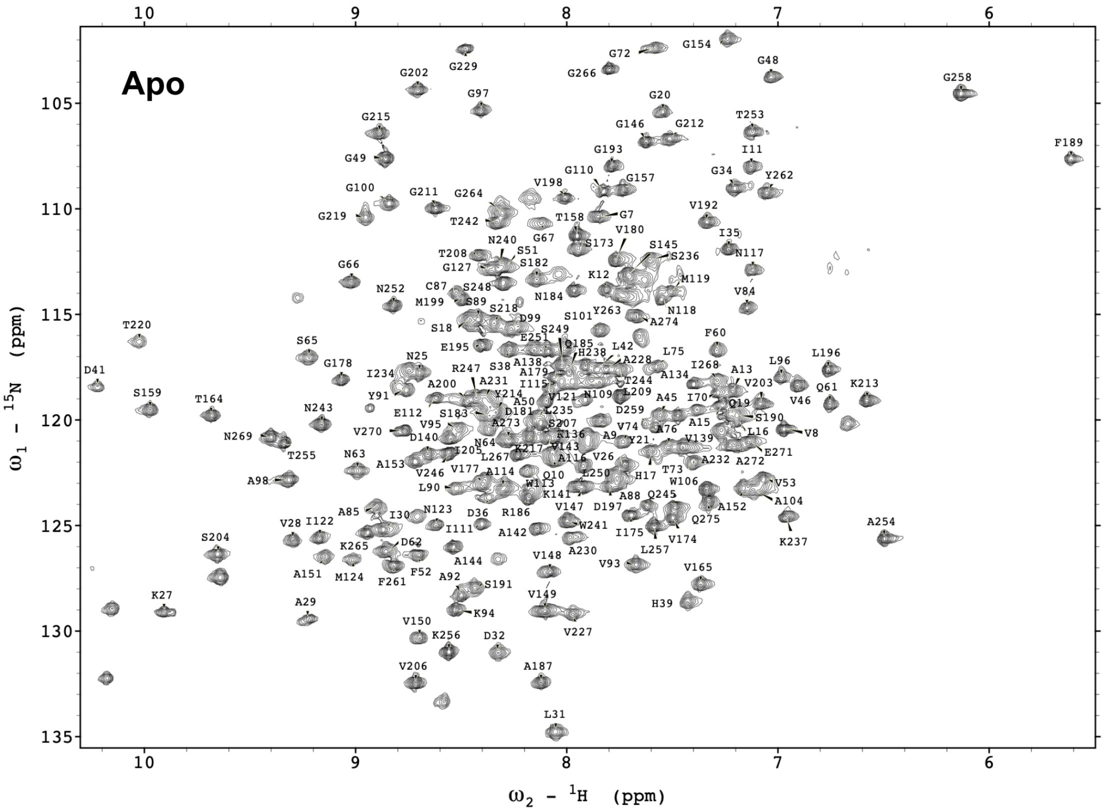


**Figure S4. Two-dimensional $^{1}H$-$^{15}N$ TROSY-HSQC spectrum of RASProtease(II) in the apo form.** Assignments are shown for backbone amide resonances.

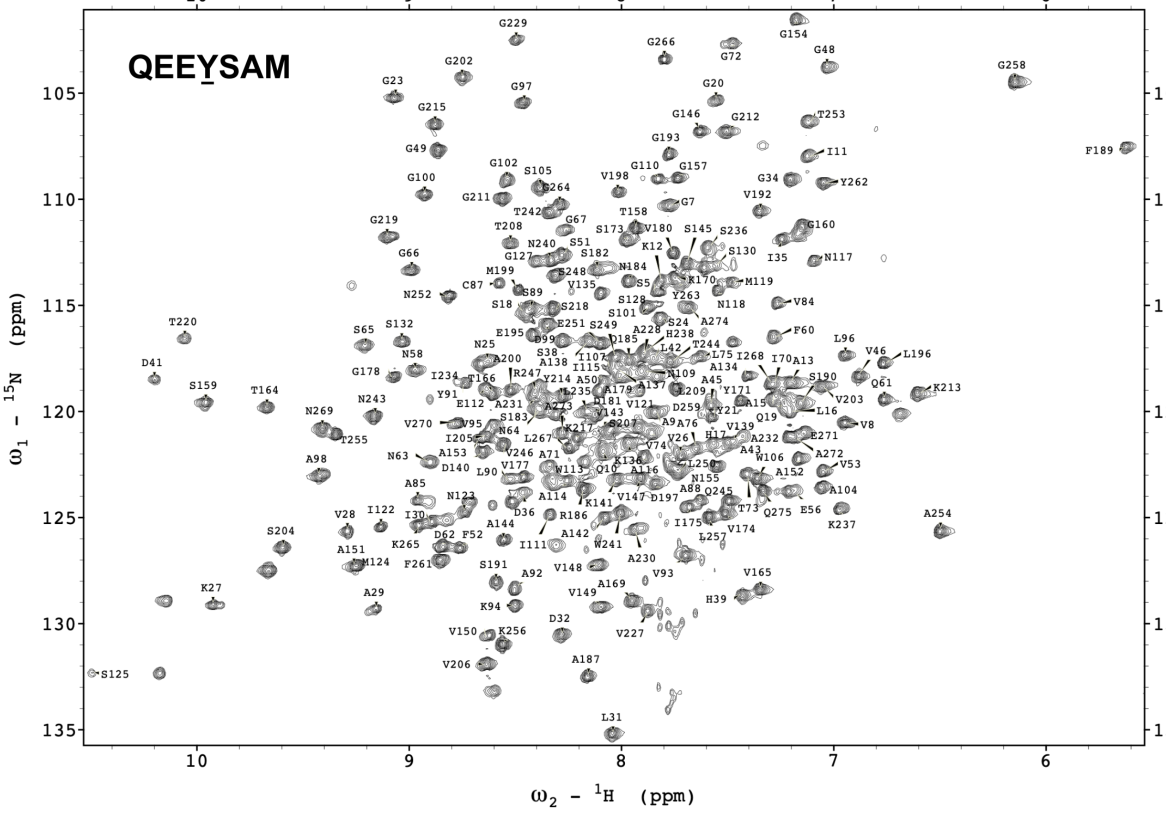


**Figure S5. Two-dimensional $^{1}H$-$^{15}N$ TROSY-HSQC spectrum of RASProtease(II) in complex with the cognate QEEYSAM peptide.** Assignments are shown for backbone amide resonances.

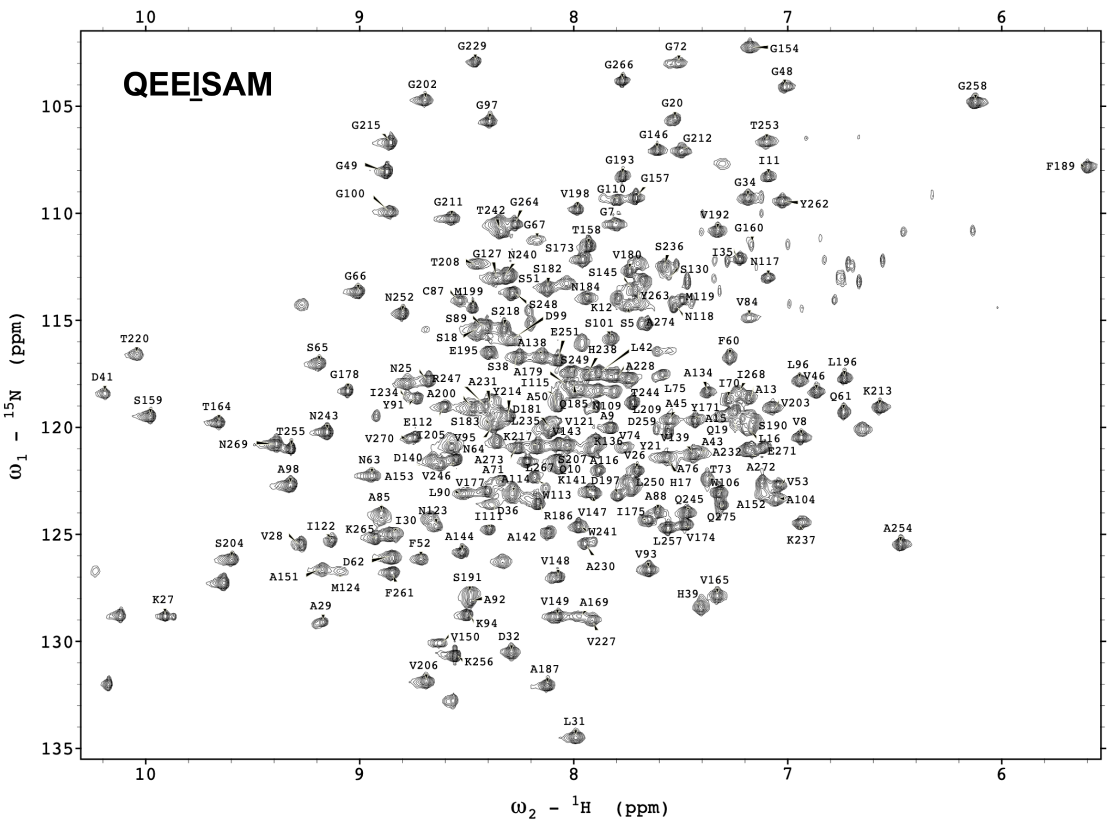


**Figure S6. Two-dimensional $^{1}$H-$^{15}$N TROSY-HSQC spectrum of RASProtease(II) in complex with the non-cognate QEEISAM peptide.** Assignments are shown for backbone amide resonances.

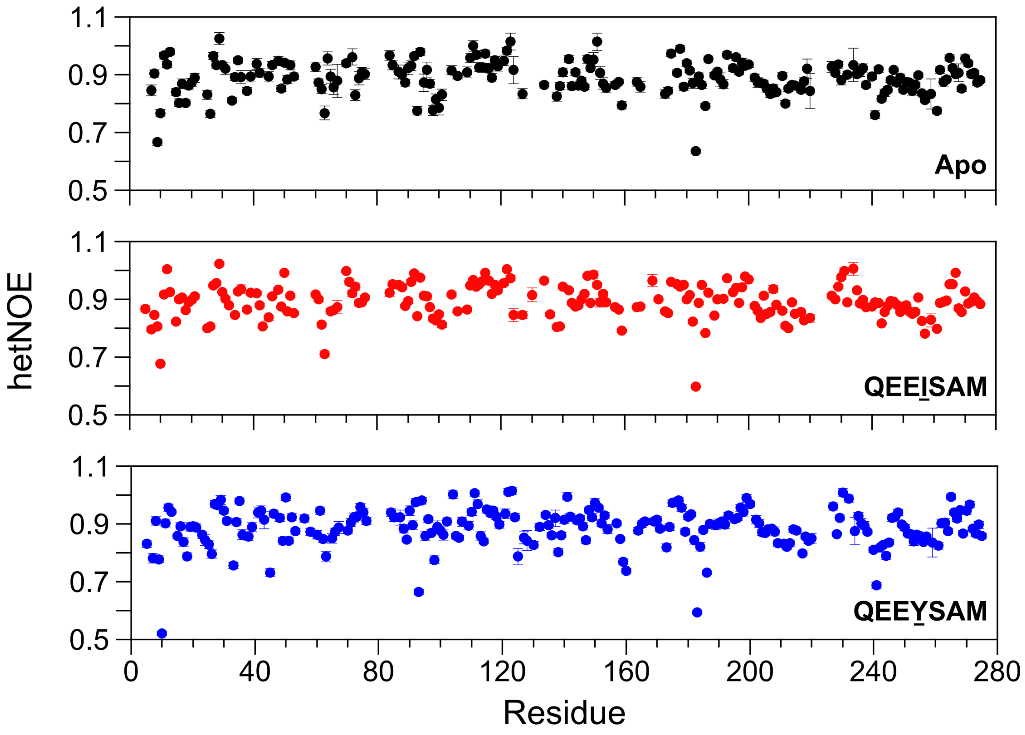


**Figure S7. Fast timescale (ps–ns) backbone amide dynamics of the apo, QEEISAM-bound, and QEEYSAM-bound RASProtease(II) forms, measured by heteronuclear NOE (hetNOE) experiments.** Individual residue steady-state {$^{1}$H}-$^{15}$N hetNOE values are plotted for RASProtease(II) in the apo form (top), in complex with QEEISAM (center), and in complex with QEEYSAM (bottom). Values greater than approximately 0.8 typically indicate restricted backbone motion of ordered regions.

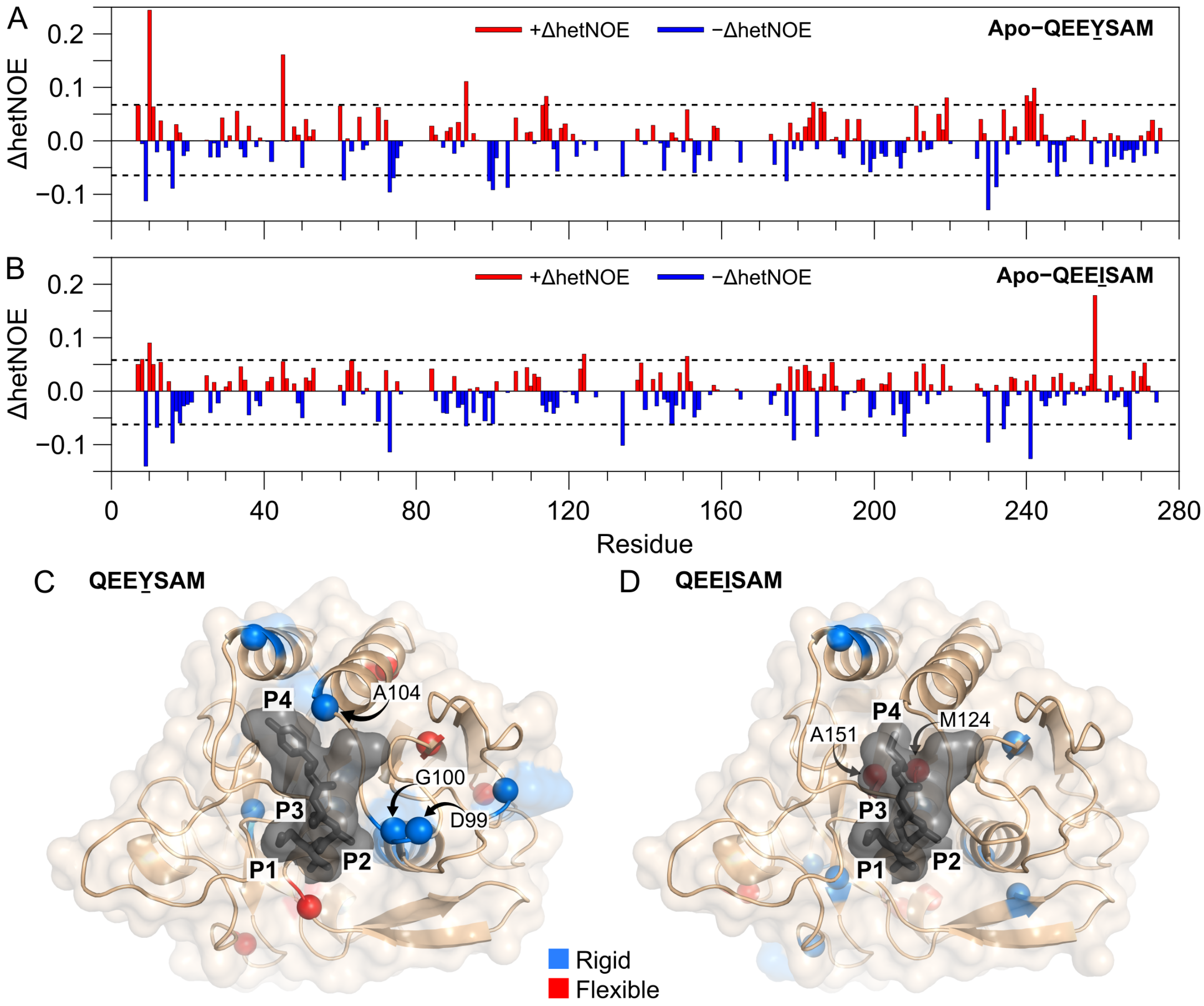


**Figure S8. Comparison of backbone amide dynamics of RASProtease(II) between the QEEYSAM-bound and QEEISAM-bound states by heteronuclear NOE (hetNOE) analysis.** **(A, B)** Differences in hetNOE values at individual residues between the apo and peptide-bound forms were calculated separately for the QEEYSAM and QEEISAM complexes. The ΔhetNOE plots show the difference between **(A)** apo and QEEYSAM and between **(B)** apo and QEEISAM. Positive ΔhetNOE values (red bars) indicate increased backbone flexibility, whereas negative ΔhetNOE values (blue bars) reflect decreased flexibility upon peptide binding. The mean ± 1.5 SD (calculated from the Apo−QEEYSAM dataset) is shown as dashed lines in each plot. This same cutoff was applied to the Apo−QEEISAM data for comparison. Residues outside this range were identified as outliers and are mapped onto the corresponding RASProtease(II) structures in complex with the **(B)** QEEYSAM or **(C)** QEEISAM peptide. Red spheres indicate increased backbone flexibility, whereas blue spheres indicate decreased flexibility upon peptide binding. The peptide is shown as gray sticks with a semi-transparent surface to highlight the active site. Only P1 through P4 residues of the peptide are shown for clarity.

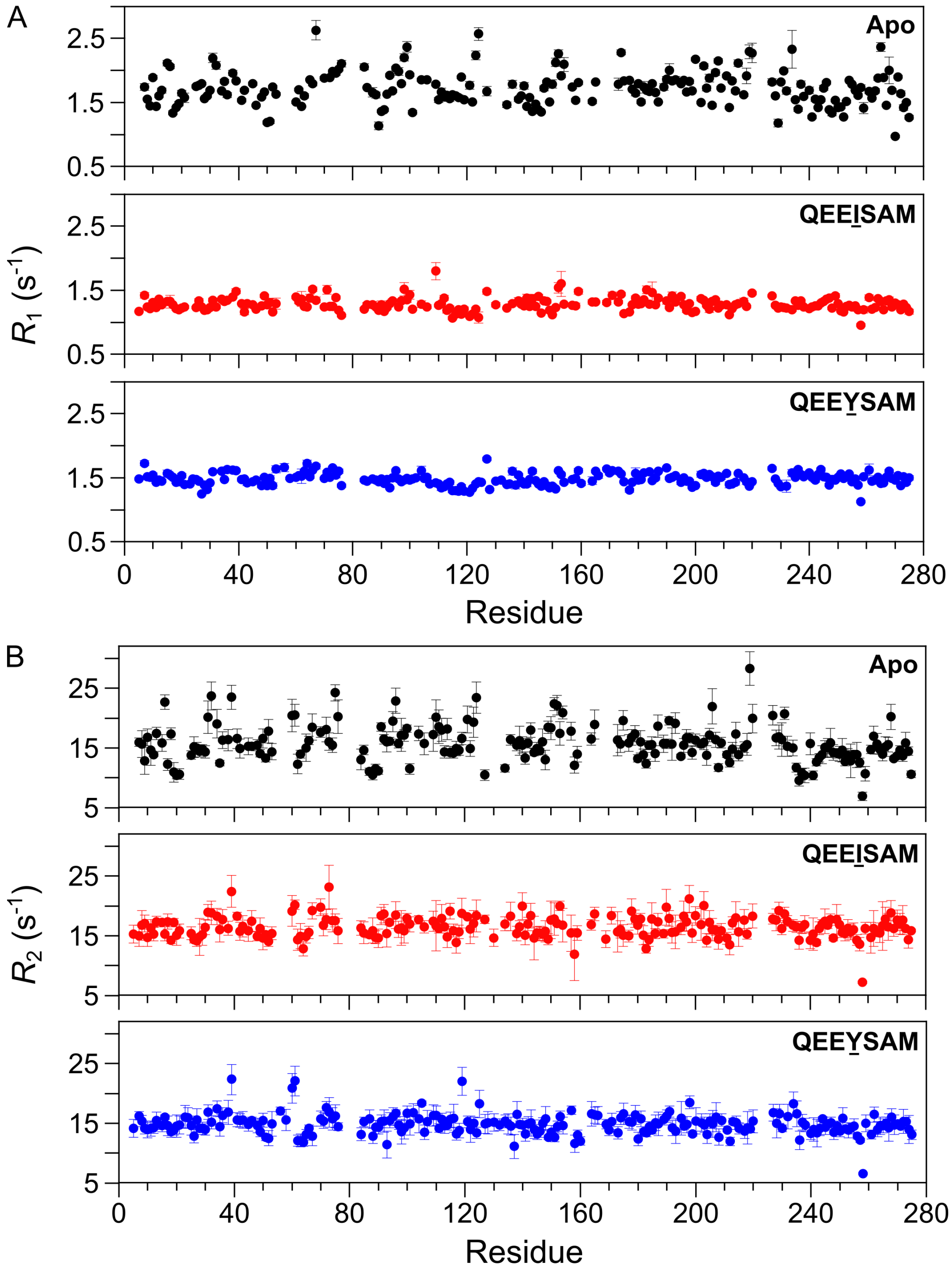


**Figure S9. Residue-specific backbone relaxation rates of RASProtease(II) in the apo form, non-cognate QEEISAM, and cognate QEEYSAM complexes. (A)** Longitudinal ($R_1$) and **(B)** transverse ($R_2$) $^{15}$N relaxation rates are plotted for the apo form (top), QEEISAM-bound (center), and QEEYSAM-bound (bottom) states. Error bars represent standard deviations from curve fitting.

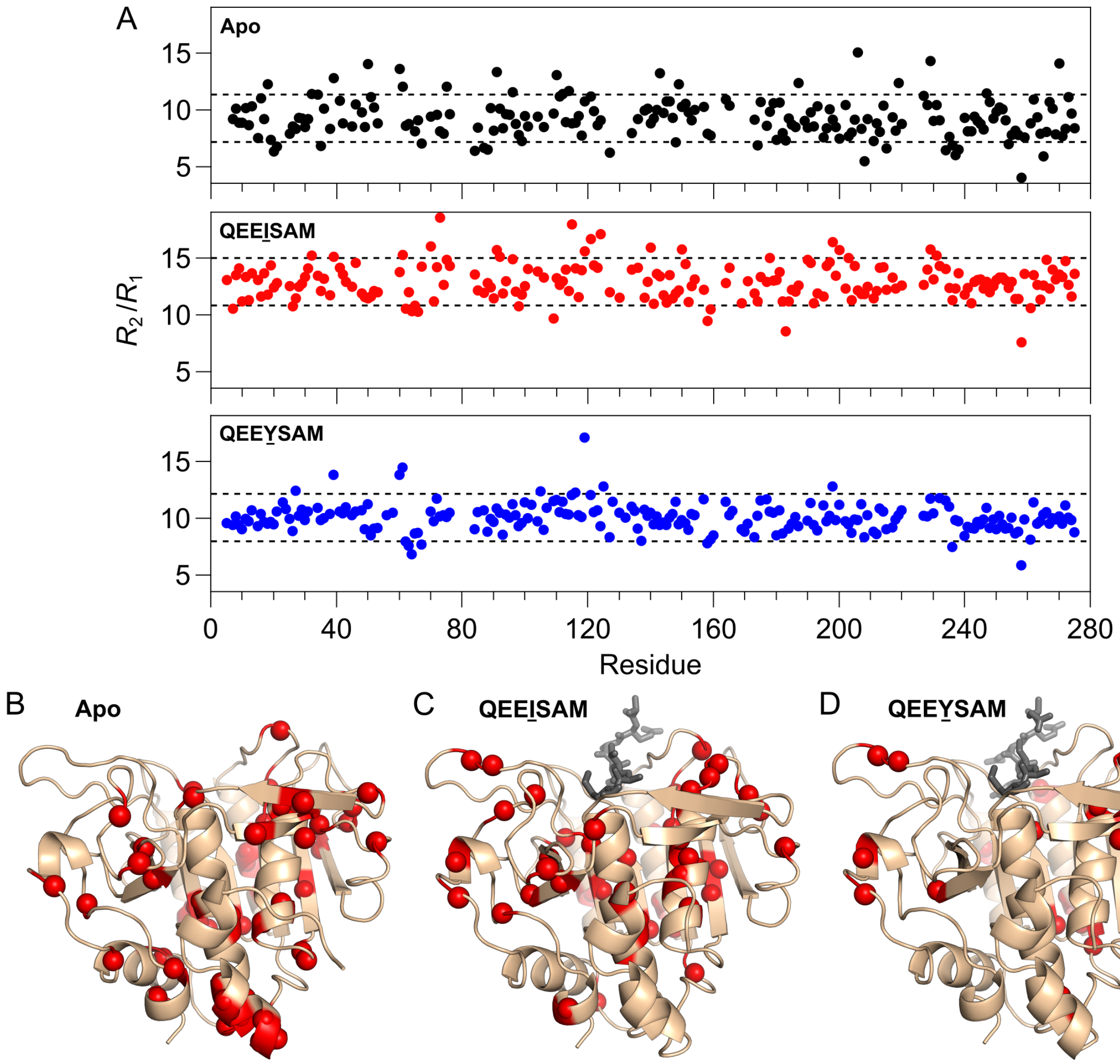


**Figure S10. Backbone conformational exchange of RASProtease(II) in the apo, QEEISAM-bound, and QEEYSAM-bound forms estimated from $R_2/R_1$ ratios. (A)** Residue-specific $R_2/R_1$ ratios were calculated for each form: apo (top), QEEISAM-bound (center), and QEEYSAM-bound (bottom). The mean ± 1 SD cutoff is shown as dashed lines in each plot. To facilitate direct comparison, the apo, QEEISAM, and QEEYSAM thresholds are defined as follows: apo mean ± apo 1 SD, QEEISAM mean ± apo 1 SD, and QEEYSAM mean ± apo 1 SD, respectively. Residues outside this range were identified as outliers and are mapped in red onto the corresponding crystal structures of the **(B)** apo RASProtease(II), **(C)** QEEISAM-bound, or **(D)** QEEYSAM-bound complexes.

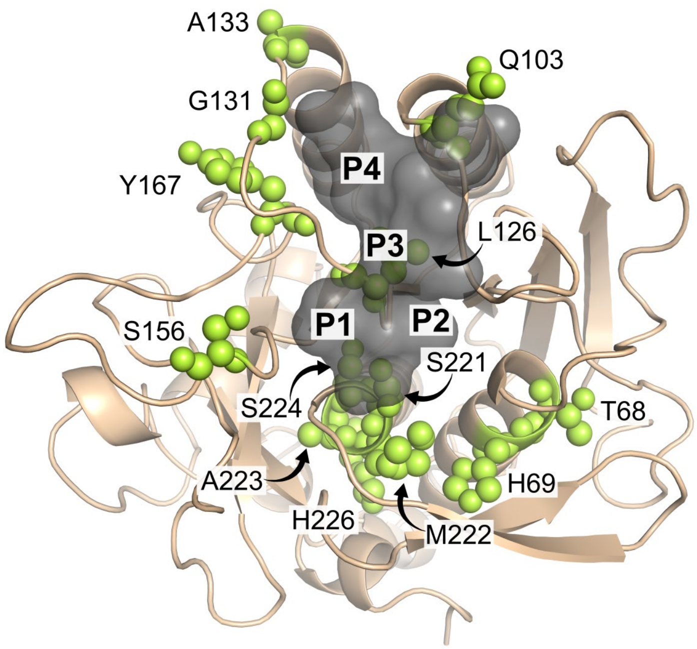


**Figure S11. A subset of putatively exchange-broadened residues common to apo, QEEISAM-bound, and QEEYSAM-bound RASProtease(II).** Residues lacking NMR assignments in all three forms and located near the S1–S4 pockets of the active site are shown as yellow-green spheres. This subset includes 13 residues (T68, H69, Q103, L126, G131, A133, S156, Y167, S221, M222, A223, S224, H226). The QEEYSAM structure is used to represent the active site, which is shown as a gray semi-transparent surface. Only P1–P4 positions of the peptide are labeled for clarity.

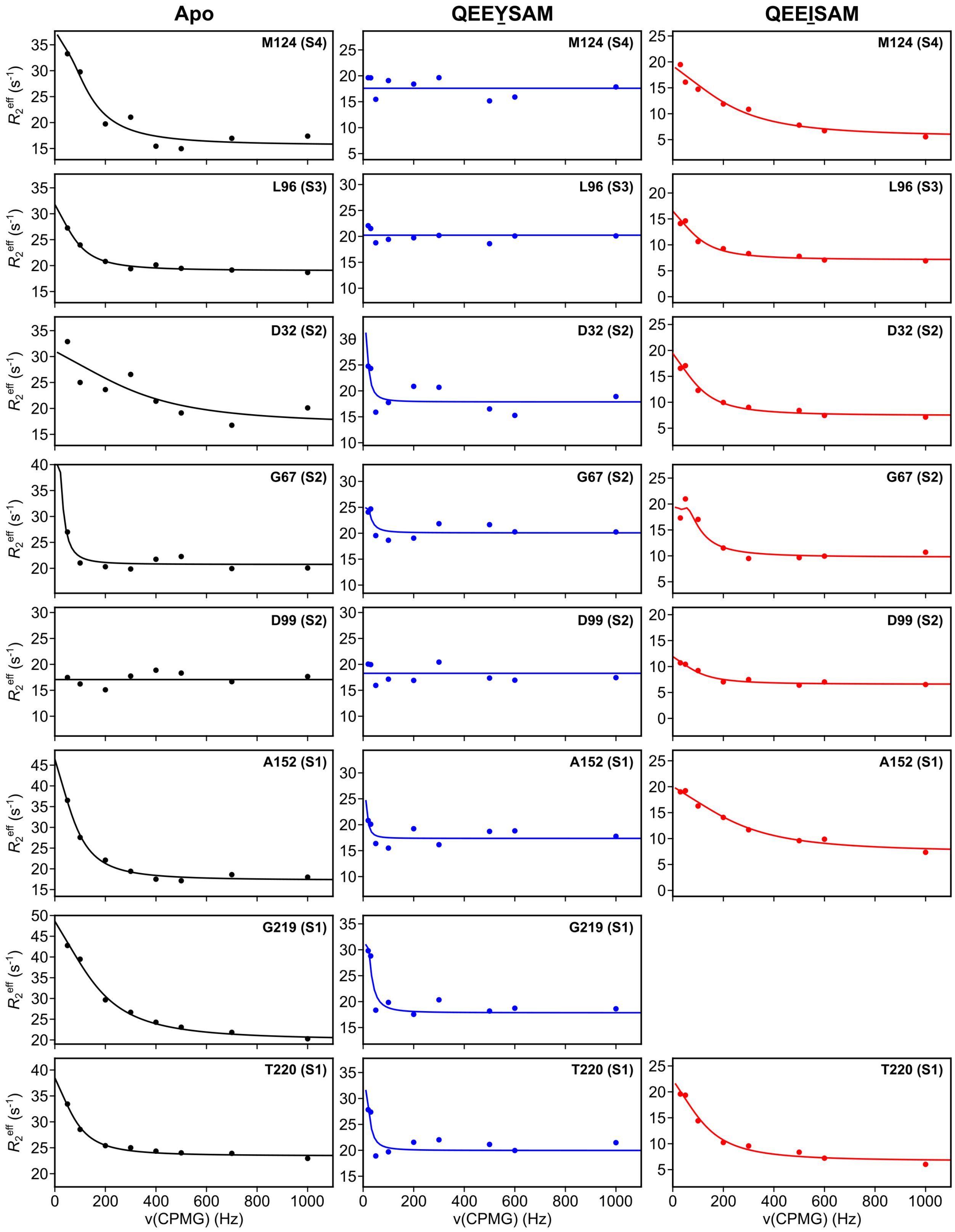


**Figure S12. Relaxation dispersion (RD) analysis of residues proximal to the active site of RASProtease(II) in the apo, QEEYSAM-bound, and QEEISAM-bound forms.** RD profiles are shown for 8 proximal residues—M124, L96, D32, G67, D99, A152, G219, and T220—for the apo form (left, black), cognate QEEYSAM complex (center, blue), and non-cognate QEEISAM complex (right, red). The nearest active site pocket (S1–S4) is noted in parentheses. Effective transverse relaxation rates ($R_{2,\text{eff}}$) are plotted as a function of CPMG field strength to probe conformational exchange on the μs–ms timescale. The curves represent fits to a two-site exchange model. Residue G219 could not be assigned in the QEEISAM-bound complex due to significant exchange broadening, so therefore its RD profile is not shown.

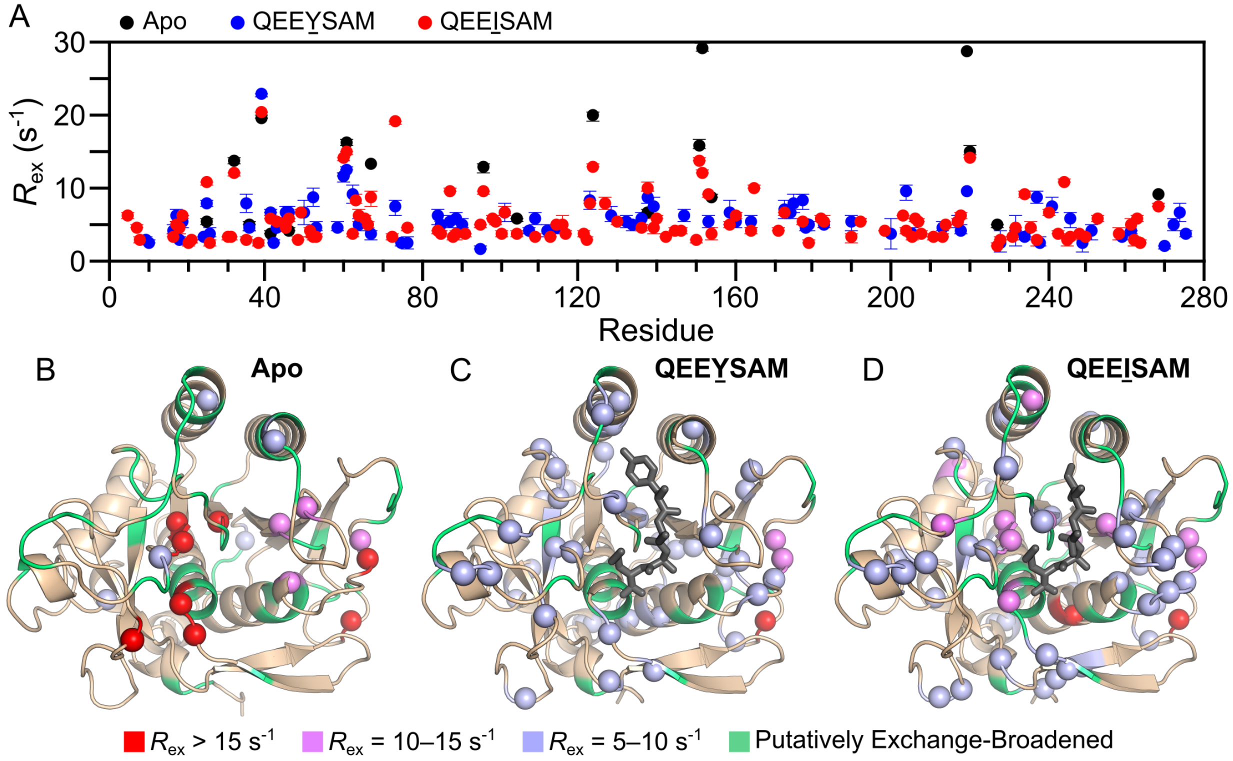


**Figure S13. Conformational exchange dynamics of RASProtease(II) induced by cognate or non-cognate peptide binding. (A)** Conformational exchange contributions ($R_{ex}$), derived from fitting of the RD data, are plotted for the apo form (black), cognate complex (blue), and non-cognate complex (red). Corresponding crystal structures of RASProtease(II) are shown for **(B)** the apo form, **(C)** QEEYSAM-bound, and **(D)** QEEISAM-bound complexes, with residues exhibiting significant conformational exchange displayed as spheres and colored according to $R_{ex}$ magnitude: red ($R_{ex}$ > 15 $s^{-1}$), violet ($R_{ex}$ = 10–15 $s^{-1}$), and light purple ($R_{ex}$ = 5–10 $s^{-1}$). Residues lacking NMR assignments are presumed to be exchange broadened and are collectively labeled "Putatively Exchange Broadened" (light green). Only P1 through P4 residues of the peptide are shown for clarity.

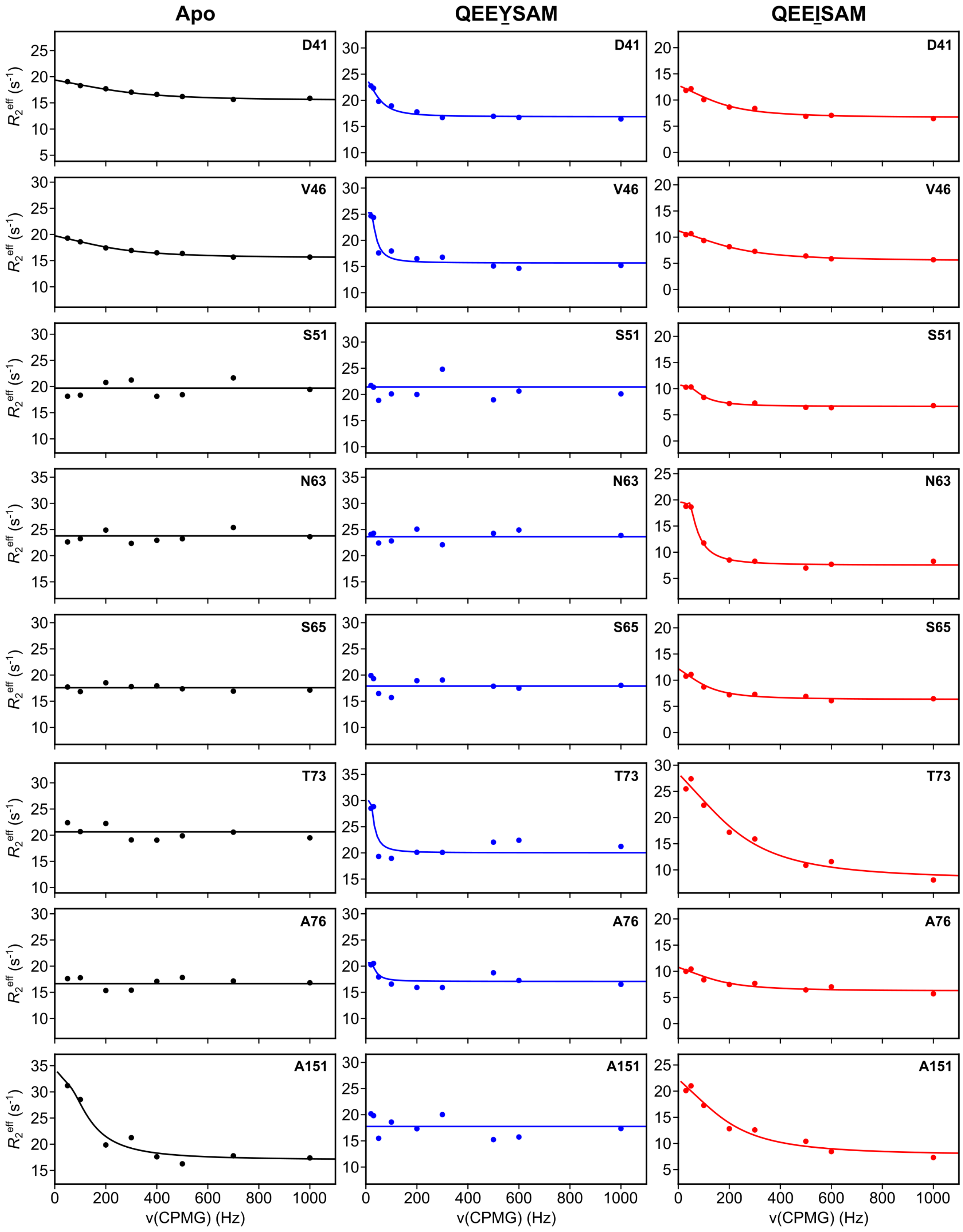


**Figure S14. Relaxation dispersion (RD) analysis of residues distal from the active site of RASProtease(II) in the apo, QEEYSAM-bound, QEEISAM-bound forms.** RD profiles are shown for 8 distal residues—D41, V46, S51, N63, S65, T73, A76, and A151—for the apo form (left), cognate QEEYSAM complex (center), and non-cognate QEEISAM complex (right). Effective transverse relaxation rates ($R_{2,\mathrm{eff}}$) are plotted as a function of CPMG field strength to probe conformational exchange on the μs–ms timescale. The curves represent fits to a two-site exchange model.

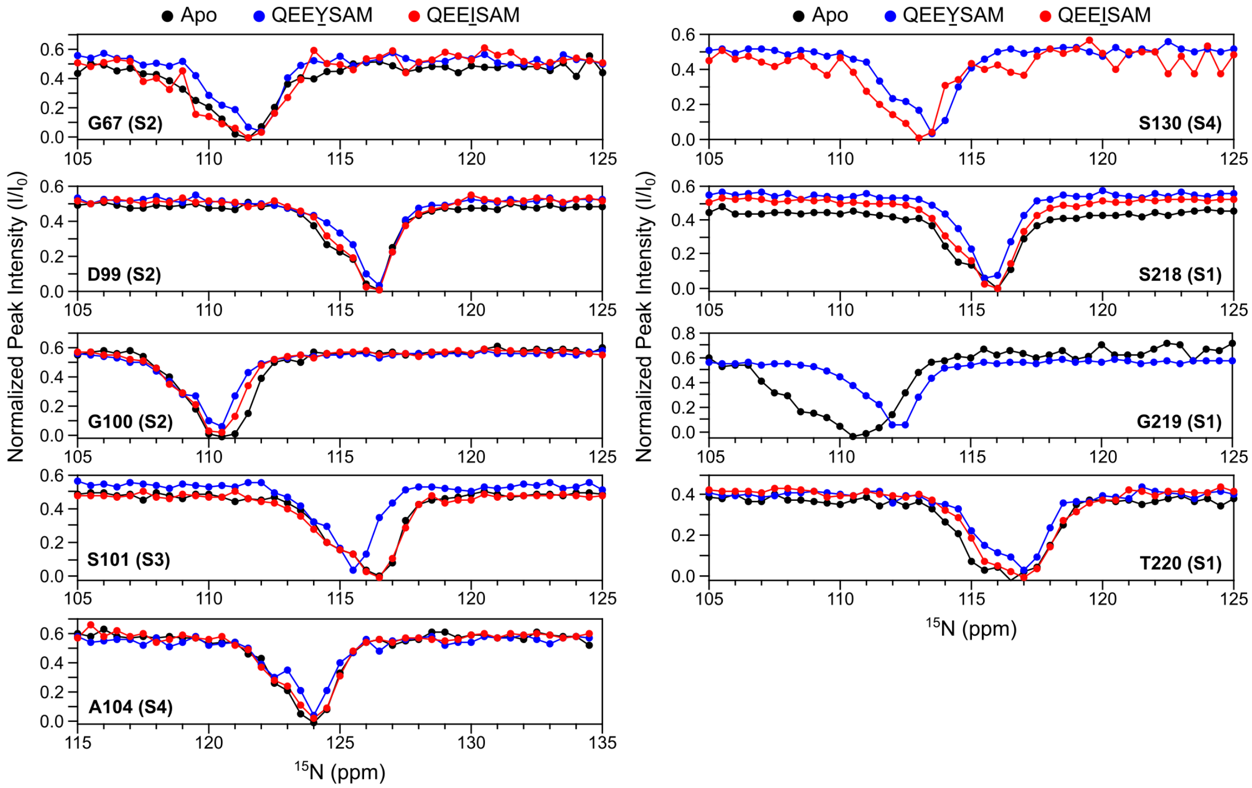


**Figure S15. Chemical exchange saturation transfer (CEST) analysis of additional residues proximal to the active site of RASProtease(II) in the apo, QEEYSAM-bound, and QEEISAM-bound forms.** CEST profiles are shown for 9 proximal residues not included in the main text—G67, D99, G100, S101, A104, S130, S218, G219, and T220—for the apo form (black), cognate QEEYSAM complex (blue), and non-cognate QEEISAM complex (red). The nearest active site pocket (S1–S4) is noted in parentheses. Normalized peak intensities are plotted as a function of $^{15}N$ saturation frequency to detect conformational exchange processes that occur on the μs–ms timescale.

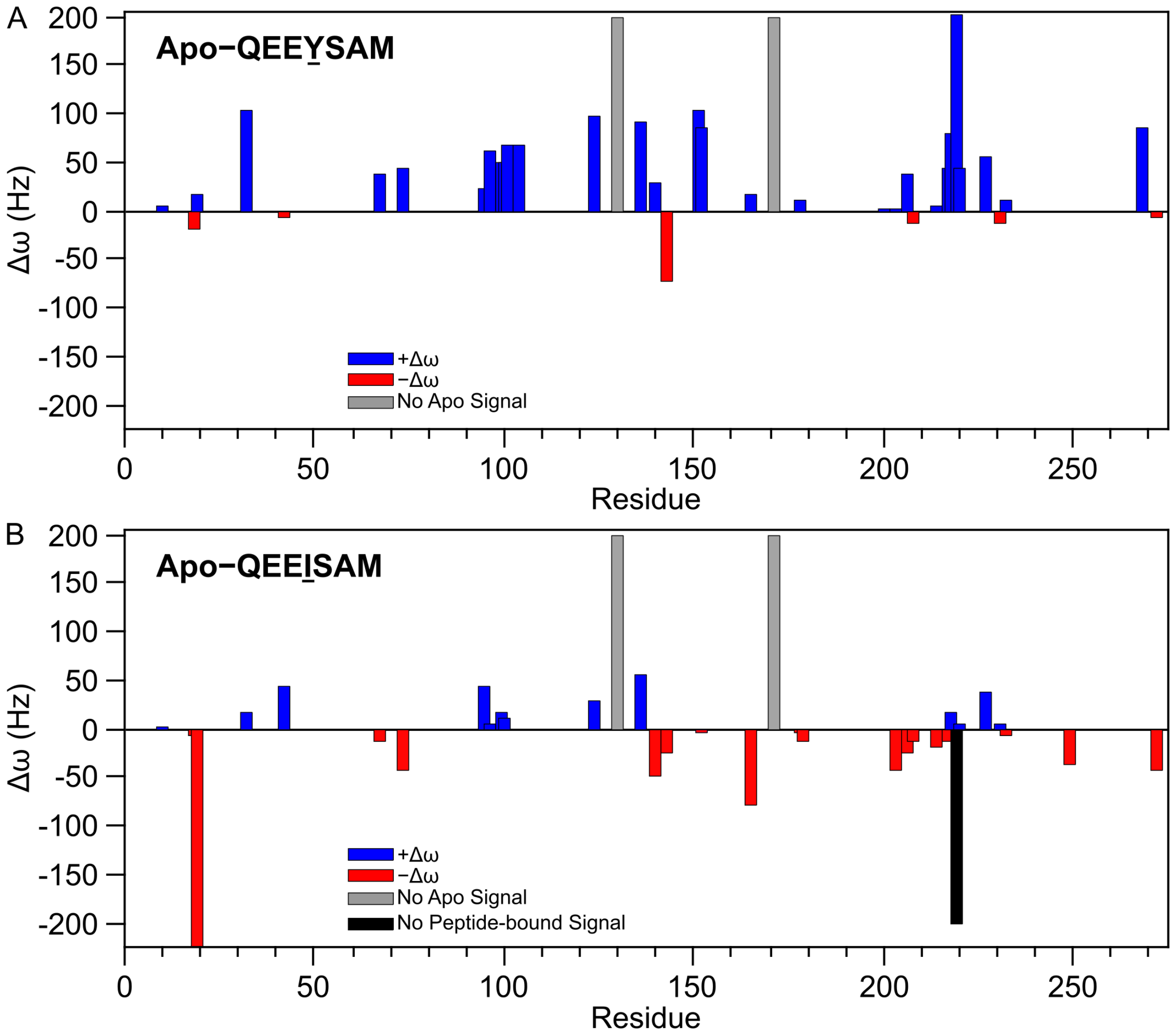


**Figure S16. Comparison of CEST profile differences (Δω) of RASProtease(II) between the QEEYSAM-bound and QEEISAM-bound states by CEST analysis.** Differences in CEST profiles (Δω) at individual residues between the apo and peptide-bound forms were calculated separately for the QEEYSAM and QEEISAM complexes. The Δω plots show the difference between **(A)** apo and QEEYSAM and between **(B)** apo and QEEISAM. Positive Δω values (blue bars) indicate line narrowing (decreased exchange), while negative Δω values (red bars) reflect line broadening (increased exchange) upon peptide binding. Gray bars indicate residues where signal was present in the peptide-bound form but missing in the apo form. The black bar represents residue G219, where signal was present in the apo form but missing in the QEEISAM-bound form. Residues lacking sufficient data are omitted.

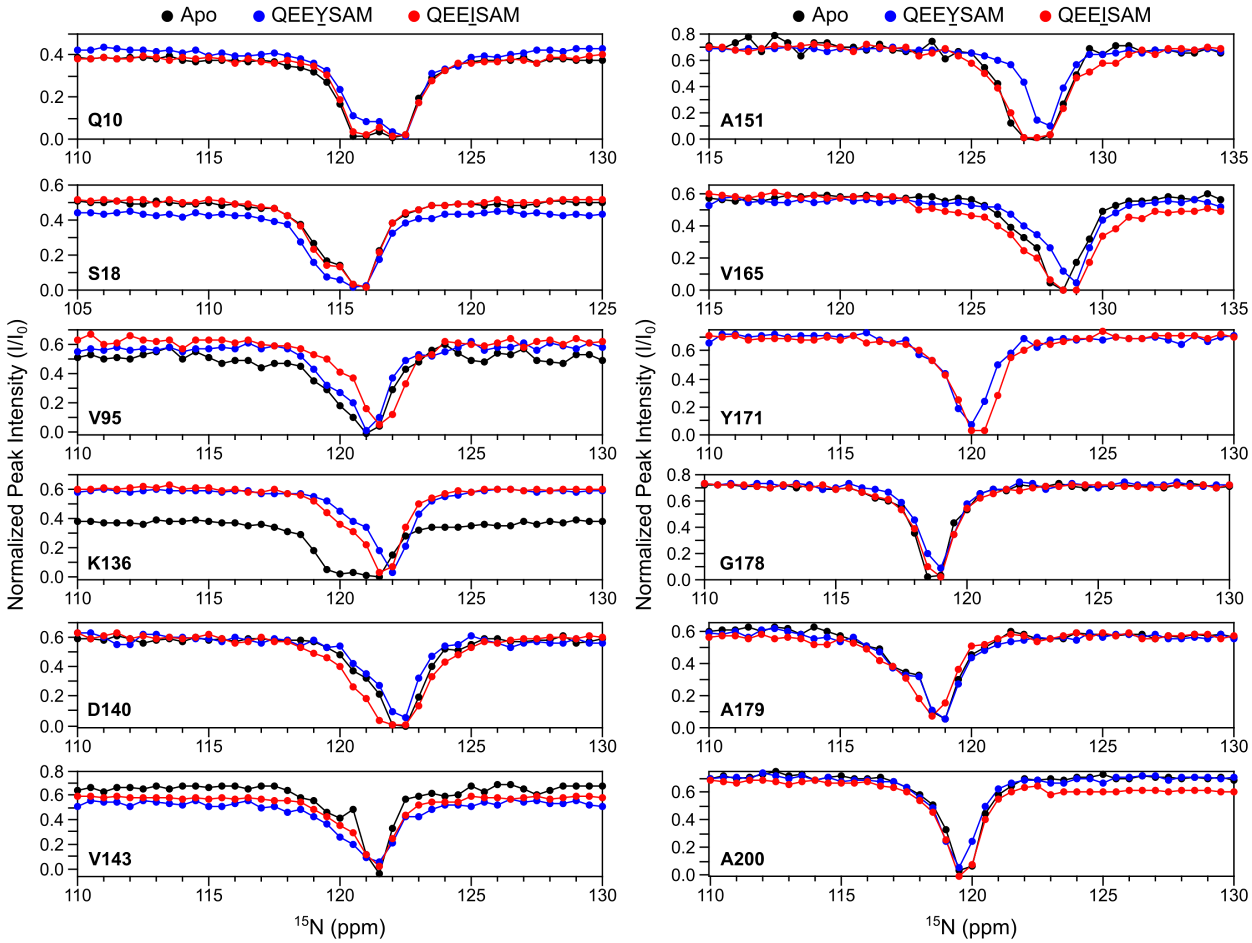


**Figure S17. Chemical exchange saturation transfer (CEST) analysis of additional residues distal from the active site of RASProtease(II) in the apo, QEEYSAM-bound, and QEEISAM-bound forms.** CEST profiles are shown for 22 distal residues not included in the main text—Q10, S18, V95, K136, D140, V143, A151, V165, Y171, G178, A179, A200, V203, V206, T208, Y214, K217, V227, A231, A232, S249, and A272—for the apo form (black), cognate QEEYSAM complex (blue), and non-cognate QEEISAM complex (red). Normalized peak intensities are plotted as a function of $^{15}N$ saturation frequency to detect conformational exchange processes that occur on the μs–ms timescale.

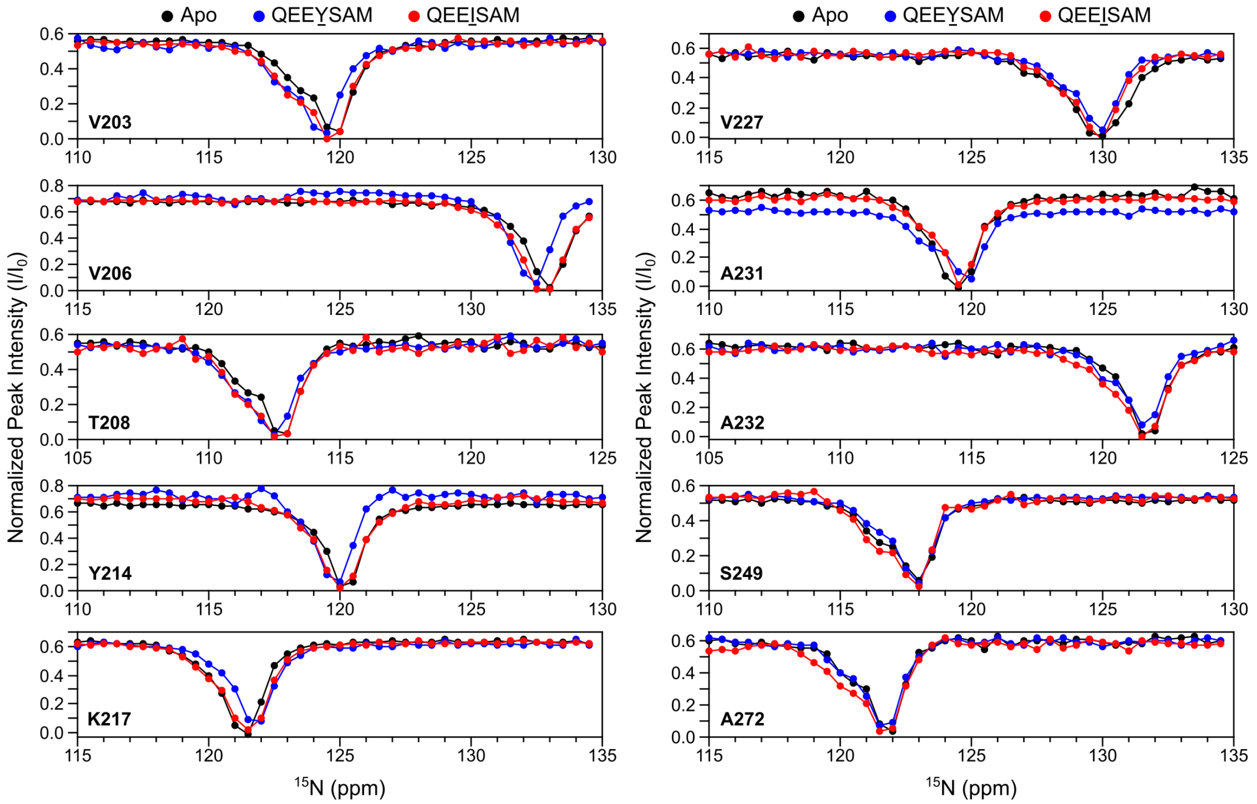


**Figure S17 (continued). Additional CEST profiles for distal residues of RASProtease(II), in the apo, QEEYSAM-bound, and QEEISAM-bound forms, as described in Figure S17.**
